# Voltage-gated sodium channel activity mediates sea urchin larval skeletal patterning through spatial regulation of Wnt5 expression

**DOI:** 10.1101/2022.11.18.517086

**Authors:** Christopher F. Thomas, Dakota Y. Hawkins, Viktoriya Skidanova, Simone R. Marrujo, Janay Gibson, Ziqing Ye, Cynthia A. Bradham

## Abstract

Defining pattern formation mechanisms during embryonic development is important for understanding the etiology of birth defects and to inform tissue engineering approaches. In this study, we used tricaine, a voltage-gated sodium channel (VGSC) inhibitor, to show that VGSC activity is required for normal skeletal patterning in *Lytechinus variegatus* sea urchin larvae. We demonstrate that tricaine-mediated patterning defects are rescued by an anesthetic-insensitive version of the VGSC LvScn5a. Expression of this channel is enriched in the ventrolateral ectoderm where it spatially overlaps with posterolaterally expressed Wnt5. We show that VGSC activity is required to spatially restrict Wnt5 expression to this ectodermal region that is adjacent and instructive to clusters of primary mesenchymal cells that initiate secretion of the larval skeleton as triradiates. Tricaine-mediated Wnt5 spatial expansion correlates with the formation of ectopic PMC clusters and triradiates. These defects are rescued by Wnt5 knock down, indicating that the spatial expansion Wnt5 is responsible for the patterning defects induced by VGSC inhibition. These results demonstrate a novel connection between bioelectrical status and the spatial control of patterning cue expression during embryonic pattern formation.

**Summary statement:** Inhibition of voltage-gated sodium channels perturbs Wnt5-mediated patterning of the sea urchin larval skeleton

## Introduction

The movement of ions is responsible for regulating processes such as wound healing, regeneration, and development (Adams and Levin, 2006; Adams et al., 2007; Akasaka et al., 1997; Beane et al., 2013; Cole and Woodruff, 2000; Hu et al., 2018; Kawakami et al., 2005; Levin, 2012, 2014; Nuccitelli, 2003; Schatzberg et al., 2015; Shipp and Hamdoun, 2012; Stumpp et al., 2012). During sea urchin development, ion channels mediate both the fast and slow blocks to polyspermy and play a role in regulating left-right symmetry breaking, ventral specification, skeletal patterning, and PMC biomineralization (Bergeron et al., 2010; Hibino et al., 2006; Jaffe, 1976; Piacentino et al., 2016a; Schatzberg et al., 2015). To uncover additional roles for ion channels in sea urchin development, we performed a screen of ion channel inhibitors which revealed that inhibition of voltage-gated sodium channels (VGSCs) by tricaine elicits larval skeletal patterning defects. Tricaine is a commonly used anesthetic and a potent inhibitor of VGSCs (Attili and Hughes, 2014; Cakir and Strauch, 2005; Frazier and Narahashi, 1975; Letcher, 1992; Schoettger, 1967).

The calcium carbonate sea urchin larval skeleton is secreted by the primary mesenchyme cells (PMCs) (Lyons et al., 2014). Prior to gastrulation, the PMCs undergo an epithelial-to-mesenchymal transition (EMT) and ingress into the blastocoel. At late gastrula stage, the PMCs have migrated into a stereotypical spatial pattern comprised of a posterior ring of cells around the hindgut with bilateral ventrolateral clusters of cells; cords of cells extend from the clusters towards the anterior territory. This cellular “ring-and-cords” organization and the corresponding skeletal elements are considered the primary (1°) pattern. Biomineralization is initiated in the clusters to produce bilateral skeletal triradiates; the radii are then extended as spicules along the 1° pattern. Subsequent PMC migration from the ring and cords produces the secondary (2°) pattern that yields the long skeletal elements (Ettensohn and Malinda, 1993; Piacentino et al., 2015; Piacentino et al., 2016b).

von Ubisch first suggested that cues from the ectoderm are responsible for organizing the PMCs within the blastocoel (von Ubisch, 1937). Transplantation experiments were key in confirming that PMC positioning is controlled by cues received from the ectoderm (Armstrong et al., 1993). Subsequent studies identified vascular endothelial growth factor (VEGF) as a patterning cue (Duloquin et al., 2007). VEGF is initially expressed in two ventrolateral patches in the ectoderm, while the expression of its receptor VEGFR is enriched in the adjacent PMC clusters (Adomako-Ankomah and Ettensohn, 2013; Duloquin et al., 2007). Wnt5 is another ventrolateral ectodermal patterning cue that is required for biomineralization and is sufficient to induce abnormal PMC migration (McIntyre et al., 2013). Our group uncovered other ectodermal patterning cues, including the sulfate transporter SLC26a2/7 (Piacentino et al., 2016b). However, the roles that other ion channels and pumps play in regulating skeletal patterning remain incompletely understood.

VGSCs induce action potentials in excitable cells such as neurons, and knockout experiments in mice show that VGSCs are required for normal development of the central nervous system (CNS) (Harris and Pollard, 1986; Planells-Cases et al., 2000; Yu et al., 2006). In zebrafish, expression of the VGSC isoform Na_v_1.5 is required for normal heart development (Chopra et al., 2010). VGSCs have also been identified in numerous other cell types including fibroblasts, glia, immune cells, and cancer cells where their functions remain less clear (Black and Waxman, 2013; Brackenbury et al., 2008; de Lera Ruiz and Kraus, 2015; Kaestner et al., 2018; Roger et al., 2015).

In this study, we show that inhibition of VGSC activity in *Lytechinus variegatus* (Lv) embryos elicits skeletal patterning defects that can be rescued via overexpression of an anesthetic insensitive version of the VGSC Scn5a. We show that VGSC activity is required during gastrulation for normal skeletal patterning and for normal spatial positioning of PMCs. We find that VGSC inhibition results in spatial expansion of Wnt5 in the ventrolateral ectoderm, the presence of ectopic PMC clusters that express Jun, and the occurrence of supernumerary skeletal triradiates. Finally, we show that tricaine-mediated skeletal patterning defects are rescued by partial knockdown of Wnt5 but not VEGF. Together, these results establish a crucial role for VGSC activity in the spatial restriction of Wnt5 in the ventrolateral ectoderm that is required for normal skeletal patterning and implicate Wnt5 as a VEGF-independent skeletal patterning cue.

## Results

### VGSC activity is required for normal skeletal patterning during gastrulation

Through a screen, we discovered that treatment with tricaine, an inhibitor of voltage-gated sodium channels (VGSCs), elicits skeletal patterning defects (Fig. 1A-B). We selected an optimal tricaine dose of 500 *µ*M as a minimal dose that results in fully penetrant skeletal patterning defects. At this dose, we did not observe changes in larval swimming behaviors, suggesting that this level of tricaine is insufficient to inhibit neural activity. We scored the defects using our established rubric (Piacentino et al., 2016b) (see Methods) and identified specific skeletal patterning abnormalities (Fig. 1B). The ventral transverse (VT) rods were the most frequently absent rods in tricainetreated embryos, whereas the most commonly observed orientation defects were anterior-posterior and left-right defects (Fig. 1B). We also observed a high frequency of ectopic rods and a lower but measurable frequency of ectopic triradiates (Fig. 1B). The occurrence of ectopic triradiates is notable in that this defect has not been previously detected with other skeletal patterning cue perturbations (Adomako-Ankomah and Ettensohn, 2013; Duloquin et al., 2007; Piacentino et al., 2016a; Piacentino et al., 2015; Piacentino et al., 2016b). To determine the temporal window during which VGSC activity is required for skeletal patterning, we performed timed treatment and washout experiments with tricaine over the time course of larval development (Fig. S1A), then assessed the resulting embryos for skeletal patterning defects at the pluteus stage (Fig. 1C-D, S1). We found that inhibiting VGSCs prior to 14 hpf results in a large proportion of embryos with patterning defects, while treatment after 14 hpf results in the majority of embryos developing control-like skeletons (Fig. 1C, S1B1-7, S1C). In the converse experiments, we found that inhibiting VGSCs from fertilization until 17 hpf results in the majority of embryos developing defective skeletons. However, removal of tricaine at 16 hpf or earlier results in normal skeletal patterns within the majority of larva (Fig. 1D, S1). We scored the patterning defects in embryos exposed to tricaine for these defined temporal intervals using broad bins and found that primary skeletal defects exhibited a saddle-like profile, dominating the earliest and latest intervals, while orientation and secondary defects were most evident in the latest intervals (Fig. S1D-E). Taken together, these data show that tricaine-sensitive VGSC activity is required during gastrulation from 14 to 17 hpf for normal skeletal patterning.

**Figure 1.**
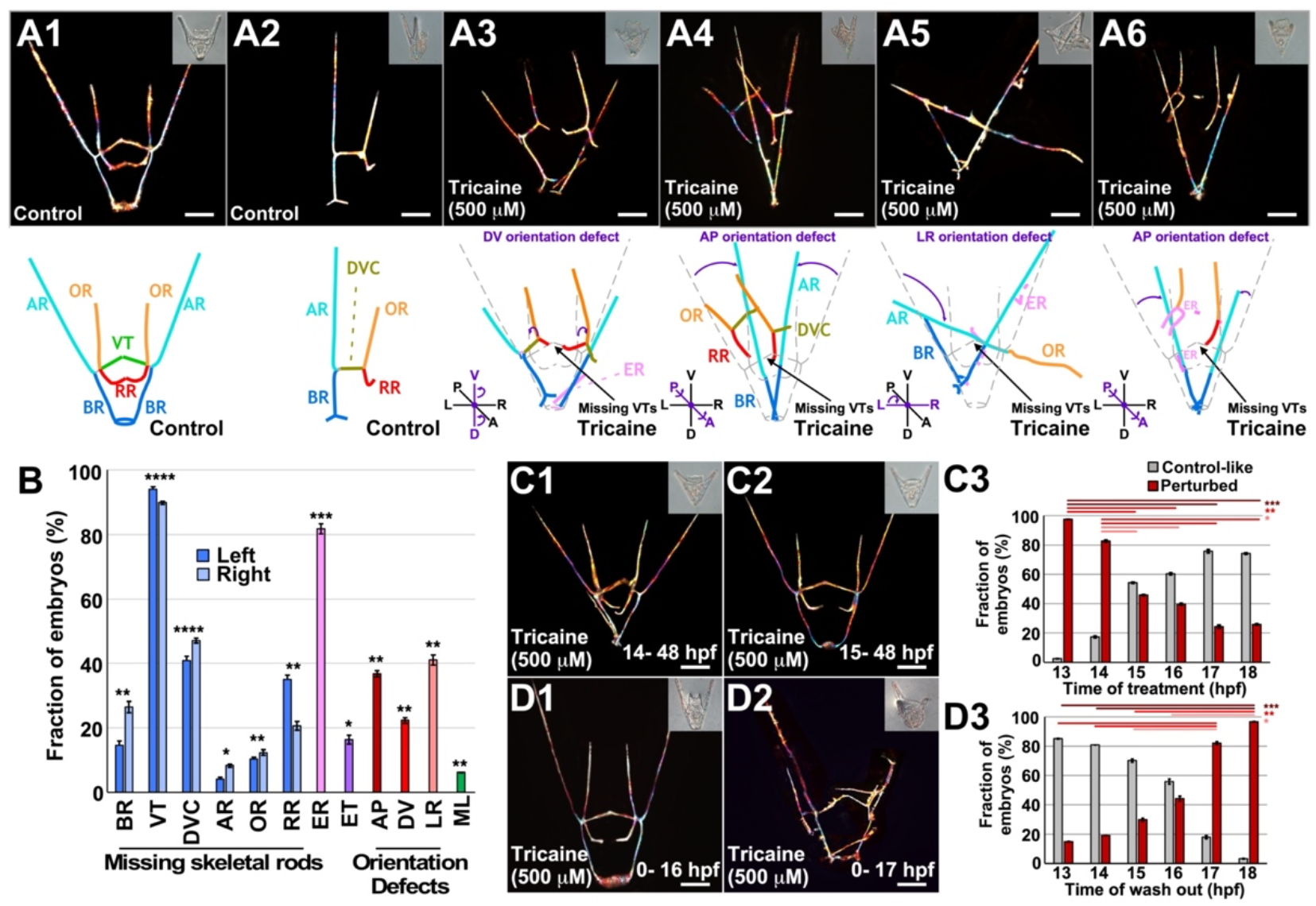
Voltage-gated sodium channel (VGSC) activity is required for normal skeletal patterning during gastrulation. A. Representative skeletal birefringence images of controls (A1-2, upper panels) and embryos that were exposed to tricaine from fertilization until analysis (A3-6, upper panels) are shown at 48 hpf with corresponding morphology images (DIC) inset. Schematized versions of each skeleton are shown (A, lower panels) with skeletal elements indicated by color, including body rods (BRs, blue), aboral rods (ARs, cyan), ventral transverse rods (VTs, green), dorsal ventral connecting rods (DVCs, dark yellow), recurrent rods (RRs, red), oral rods (ORs, orange). Ectopic rods (ERs, pink) are also indicated. Orientation defect exemplars show abnormal orientations about the AP, DV, or LR axes (A3-6). B. Skeletal defects including element losses, ectopic rods (ER), ectopic triradiates (ET), orientation defects, and midline losses (ML) are plotted as the average percentage of embryos exhibiting each defect ± s.e.m.; * p < 0.05, **, p < 0.005, *** p < 0.0005, **** p < 10^−7^ compared to control (t-test); n = 49. C-D. Embryos that were treated with tricaine during the indicated hours post-fertilization (hpf) (C1, C2, D1, D2), are shown at pluteus stage (48 hpf) as skeletons (birefringence) with corresponding DIC images inset. Embryos were treated with tricaine at the indicated time points (C), or treated with tricaine at fertilization and removed from the drug at the indicated time points (D), then scored for patterning defects at pluteus stage (C3, D3); the results are shown as the average percentage of control-like and perturbed embryos ± s.e.m.; * p < 0.05, **, p < 0.005, *** p < 0.0007 (Tukey-Kramer test); n ≥ 247 (C3) or n ≥ 177 (D3) per condition. See also Fig. S1. Scale bars are 50 *µ*m.

### VGSC inhibition is sufficient to perturb both intracellular sodium ion concentration and transmembrane voltage

To functionally confirm that tricaine inhibits VGSCs in sea urchin embryos, we investigated the effect of tricaine treatment on transmembrane voltage (V_mem_) and sodium ion (Na^+^) levels by using the fluorescent reporters DiSBAC and CoroNa-AM, respectively (Adams and Levin, 2012; Epps et al., 1994; Meier et al., 2006; Rodriguez-Sastre et al., 2019). To validate tricaine, we first subjected the embryos to tricaine treatment only minutes prior to measuring V_mem_ and Na^+^ levels to determine the acute response to the inhibitor. We found that acute VGSC inhibition results in significantly reduced Na^+^ levels relative to controls, as expected, both globally and within each of the ectodermal regions measured, including posterior ventrolateral tissue adjacent to the PMC clusters, the apical territory, and the intervening anterolateral ectodermal region (Fig. 2A1-2, C). When we similarly measured V_mem_ in the ectoderm, we found that tricaine-treated embryos are significantly hyperpolarized relative to controls, reflected as decreased depolarization globally as well as in each measured ectodermal region (Fig. 2B1-2, D), as expected upon inhibition of VGSCs. These data show that tricaine inhibits VGSC activity in sea urchin embryos, validating it functionally as a sodium channel inhibitor.

**Figure 2.**
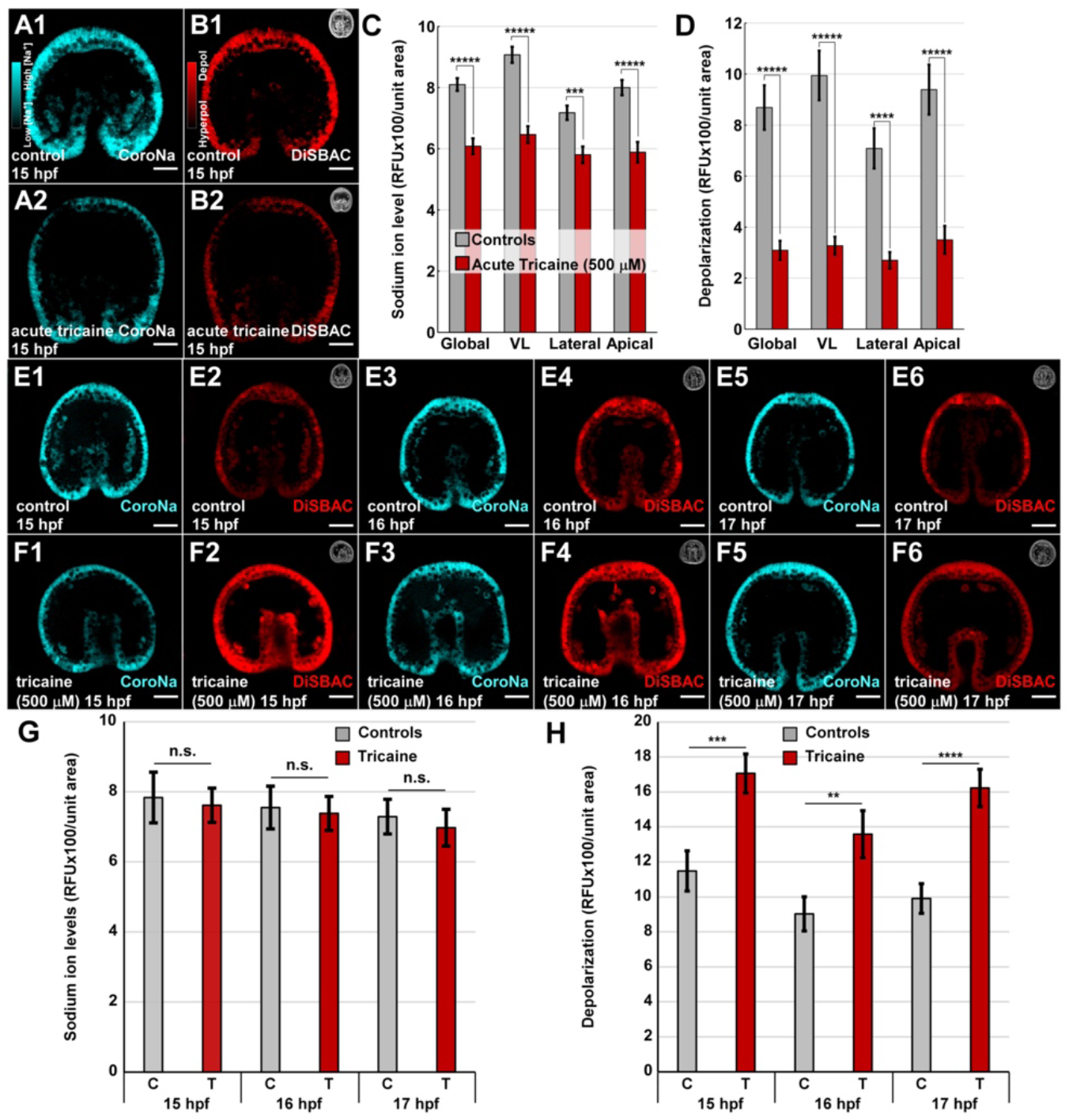
VGSC inhibition is sufficient to perturb intracellular Na^+^ concentration and membrane voltage (V_mem_). A-D. Sodium ions (Na^+^) were visualized with the fluorescent reporter CoroNa (cyan), and V_mem_ was visualized with DiSBAC (red) in the same live embryos; the signal ranges are shown by custom LUTs (A1, B1 insets). Control (A1, B1) and tricaine-treated (A2, B2) embryos were analyzed at 15 hpf immediately following tricaine treatment to determine tricaine’s acute effects. Relative Na^+^ levels (C) and V_mem_ (D) were quantified in the indicated ectodermal territories and are shown as the average RFU/area ± s.e.m.; n = 37 embryos per condition from three biological replicates; *** p < 10^−3^; ***** p < 10^−5^ (student *t*-test). E-H. Na^+^ and V_mem_ were visualized in controls (E) and embryos treated with tricaine from fertilization (F), then analyzed at the indicated time points during gastrulation. Relative signal levels were quantified as above (G-H, see also Fig. S2); n ≥ 24 per condition from two biological replicates; ** p < 10^−2^; *** p < 10^−3^; **** p < 10^−4^ (*t*-test). Scale bars represent 20 *µ*m.

Since VGSC activity is required for skeletal patterning from 14 to 17 hpf, we next asked how tricaine treatment affects V_mem_ and Na^+^ levels during this developmental window. We treated embryos with tricaine from fertilization to either mid- or late gastrula stage, corresponding to these timepoints, then measured V_mem_ and Na^+^ levels. In this case, long-term tricaine treatment did not result in a significant decrease to Na^+^ levels relative to controls (Fig. 2E, G, S2A). Instead, long-term VGSC inhibition resulted in significant depolarization of embryos relative to controls at 15, 16, and 17 hpf (Fig. 2F, H, S2B), in contrast to the acute effects of tricaine. At 7 hpf, Na^+^ levels within tricaine-treated embryos remain reduced, but the embryo has become depolarized, indicative of ion flux (Fig. S3A-C). At 11 hpf, Na^+^ levels are restored to normal levels, while depolarization persists in tricaine-treated embryos. These findings suggest that compensatory processes restore normal Na^+^ levels at the expense of V_mem_, via movement of ions, including Na^+^, through other channels, and indicate that this process is slow, requiring more than seven hours to reach homeostasis. These data also show that long-term tricaine treatment is sufficient to significantly depolarize ectodermal V_mem_ in the relevant developmental stages, while Na^+^ levels are restored to normal during this interval.

### Expression of a drug-resistant VGSC is sufficient to rescue tricaine-mediated skeletal patterning defects

VGSCs are integral membrane proteins made up of a pore-forming α subunit with associated auxiliary β subunits that enable the influx of Na^+^ into cells in response to membrane depolarization (Catterall, 2000, 2014; Ragsdale et al., 1994). The ß subunits enhance and modulate the α subunit’s function, but the α subunit can function independently of ß accessories (Wang et al., 2017; Yu et al., 2005). The ß subunits have only recently evolved and are considered a vertebrate innovation (Wang et al., 2017; Yu et al., 2005). In keeping with that, ß subunit-encoding genes have not been identified in Lv or Sp sea urchin transcriptomes (Hogan et al., 2020; Tu et al., 2014).

VGSCs are well-known mediators of neural action potentials but are also broadly expressed in non-neuronal cell types in which they appear to provide a house-keeping function (Black and Waxman, 2013; de Lera Ruiz and Kraus, 2015; Kaestner et al., 2018; Roger et al., 2015). LvScn5a is the sole annotated VGSC within the developmental transcriptome for Lv (Hogan et al., 2020). LvScn5a expression is elevated during gastrulation when embryos are sensitive to tricaine, then becomes highly elevated after gastrulation, likely corresponding with neural development (Hogan et al., 2020) (Fig. 3A). Spatial expression analysis shows that, prior to gastrulation, Scn5a expression is spatially uniform and at low levels (not shown). LvScn5a expression is enriched in the ventrolateral and apical ectoderm at mid and late gastrula stages (Fig. 3B), during the interval of sensitivity to tricaine (Fig. 1C-D), along with a lower general background level in most or all cells (Fig. 3A, S4). Between 18 and 27 hpf, the enriched expression domains of LvScn5a expand to connect the VL and apical expression domains (Fig. 3B), corresponding spatially with the ciliary band (see Fig. S6C). Since the ciliary band includes neurons, this late expression pattern matches expectations.

**Figure 3.**
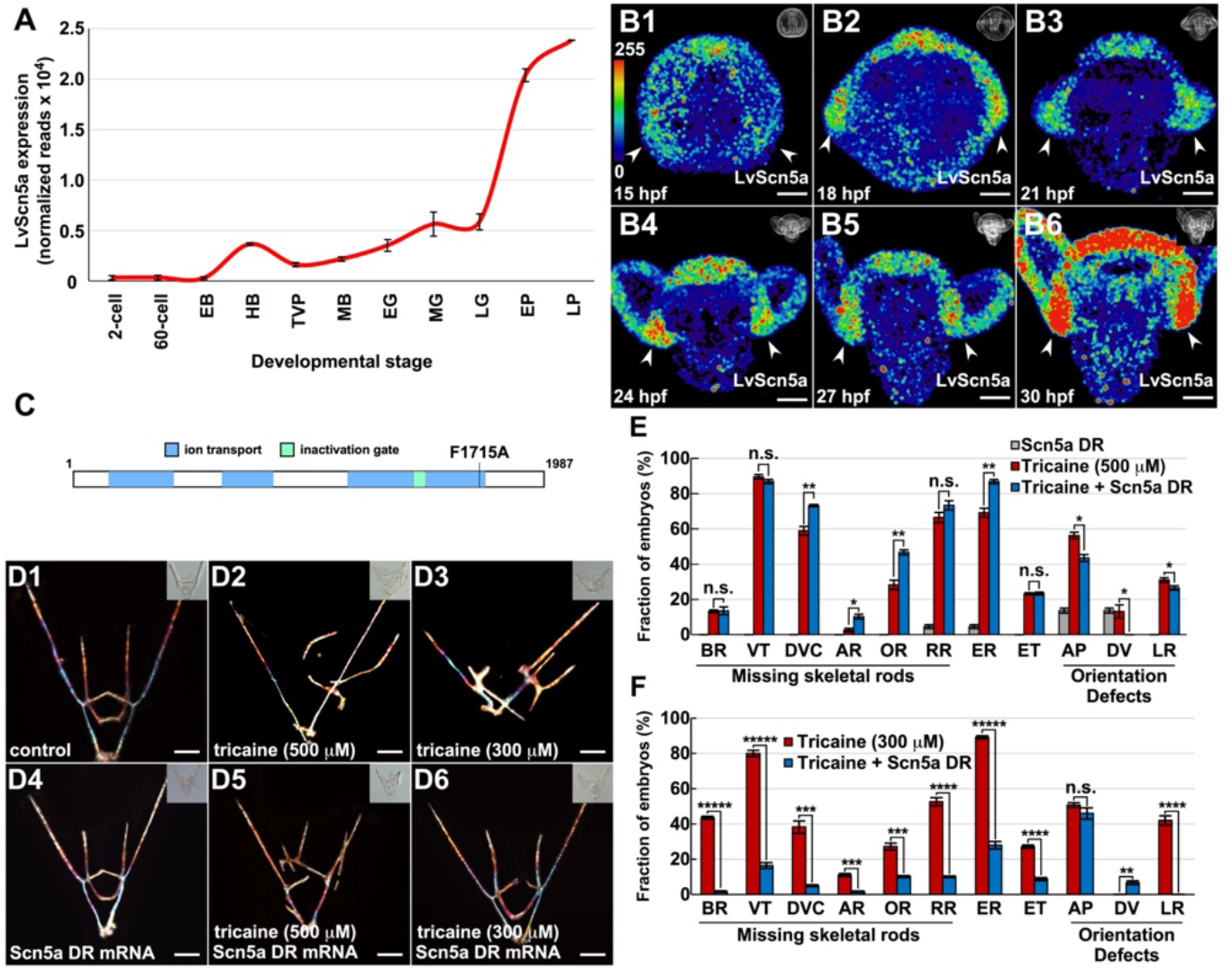
Tricaine-mediated skeletal patterning defects are rescued by overexpression of drug-resistant (DR) LvScn5a. A. The temporal expression of LvScn5a is shown at the indicated stages as normalized reads ± s.e.m. from our transcriptome sequence data (Hogan, et al., 2020). B. Control embryos were fixed at the indicated timepoints, then subjected to FISH for LvScn5a. Signal ranges in z-projected embryos are shown with a custom LUT (B1 inset). C. The schematic shows the LvScn5A gene to scale with ion transport domains and the DR mutation indicated. D. Control zygotes (D1-3) or zygotes injected with LvScn5a DR mRNA (D4-6) were untreated (D1, 4), or treated with tricaine at 500 *µ*M (D2, 5) or 300 *µ*M (D3, 6), and are shown at the pluteus stage as in Fig. 1A. The frequency of the indicated skeletal patterning defects is shown as the average percentage of embryos with each defect ± s.e.m. with tricaine at 500 (E) or 300 (F) *µ*M; see Fig. 1 for element names and abbreviations; n ≥ 30 per condition; * p < 0.05; ** p < 10^−5^; *** p < 10^−10^; **** p < 10^−20^; ***** p < 10^−40^; n.s. not significant (weighted *t*-test). Scale bars represent 50 *µ*m.

We used an LvScn5a-specific morpholino antisense oligo to knockdown Scn5a expression in an effort to phenocopy the effects of tricaine treatment. However, these experiments were unsuccessful since the resulting embryos were either normal or dead. Despite considerable optimization efforts, we were unable to identify a survivable dose that produced a phenotype. While the explanation for this outcome remains unclear, it suggests that Scn5a serves an indispensable housekeeping function, in keeping with the findings in other organisms (Black and Waxman, 2013; de Lera Ruiz and Kraus, 2015; Kaestner et al., 2018; Roger et al., 2015). We therefore turned to LvScn5a mutations as an alternative approach to test the specificity of tricaine for Scn5a.

Tricaine is a local anesthetic (LA) that reversibly inhibits VGSCs by binding to anesthetic binding sites (Cakir and Strauch, 2005; Ragsdale et al., 1994; Ramlochansingh et al., 2014). The LA binding domain in the pore of the VGSC was identified through site-directed scanning mutagenesis, then testing Na^+^ conductance (Ragsdale et al., 1994); in this way, the LA-insensitive F1760A mutant was identified. The F1760A substitution eliminates a critical binding domain for LAs, which significantly attenuates channel inhibition by LAs including tricaine (Carboni et al., 2005; Nau and Wang, 2004; Nau et al., 2000; Pless et al., 2011; Ragsdale et al., 1994; Stoetzer et al., 2016; Wright et al., 1998).

We next asked whether expression of an anesthetic-insensitive version of LvScn5a could rescue the skeletal patterning defects induced by tricaine treatment. We produced a drug-resistant (DR) version of LvScn5a by introducing the heterologous point mutation F1715A in domain IV of LvScn5a that corresponds to the F1760A mutation in human Scn5a (Fig. 3C). Overexpressed wild-type or drug resistant Scn5a is not expected to produce a phenotype since the channel requires a regulatory stimulus, voltage, to open it. We thus selected the maximum dose of Scn5a DR that remained non-phenotypic; at that dose, embryos displayed very few skeletal patterning defects, the most prevalent of which was an AP orientation defect in a small fraction of embryos (Fig. 3D4, E). The combination of DR Scn5a and 500 *µ*M tricaine resulted in skeletons that more closely approach normal patterns (Fig. 3D5); however, DR Scn5a-expressing embryos failed to exhibit a significant rescue of tricaine-induced skeletal patterning defects (Fig. 3E). The improved appearance of some tricaine-treated embryos that express DR Scn5a compared to tricaine-treated controls prompted us to repeat this rescue experiment with a lower dose of tricaine. We originally selected 500 μM as the operating dose for tricaine since the penetrance of the phenotype was nearly 100% at that dose (Fig. S5). For these experiments, we tested 300 μM, which produces the same phenotype as 500 μM, but with only 80% penetrance (Fig. S5). Controls treated with 300 μM tricaine exhibited the hallmark skeletal patterning defects associated with tricaine treatment, including loss of VTs, LR orientation defects, and the appearance of ectopic skeletal rods (Fig. 3D3, F). Embryos injected with LvScn5a DR mRNA and treated with 300 μM tricaine exhibited a dramatic and significant recovery of skeletal patterning defects, including a rescue of VTs, reduction in the frequency of ectopic rods, and no incidences of LR orientation defects (Fig. 3D6, F). The finding that LvScn5a DR mRNA is sufficient to rescue the majority of the tricaine-induced skeletal patterning defects in embryos treated with 300 μM tricaine indicates that, aside from abnormal AP orientations, the tricaine-mediated defects are due to the specific inhibition of Scn5a. Interestingly, AP orientation defects were not rescued, suggesting that tricaine treatment elicits this specific skeletal patterning defect via a mechanism other than inhibition of VGSCs. Our results suggest that the DR channel, although globally expressed, is activated in the same spatial region as the endogenous channel by endogenous voltage differences, and further imply that normal channel opening allows a maximal movement of Na^+^ since the additional open channels contributed by Scn5a DR do not produce additional phenotypic effects. The failure of LvScn5a DR mRNA to significantly rescue skeletal patterning at a higher tricaine dose suggests that the mutation introduced in the DR LvScn5a gene is insufficient to completely prevent the binding of tricaine at that dose. Therefore, at high enough tricaine concentrations, a sufficient amount of tricaine is able to bind to the exogenously expressed LvScn5a channel, resulting in its inhibition with phenotypic consequences.

### VGSC activity is not required for DV axis specification or neuronal differentiation

The observation that VGSC inhibition elicits skeletal patterning defects prompted us to assess the effects of tricaine exposure on the ectoderm. We first assessed neuronal differentiation by visualizing serotonin- and synaptotagmin B (synB)-positive neurons (Bradham et al., 2009; Yaguchi et al., 2010). Control embryos possessed both sets of neurons in normal spatial patterns (Fig. S6A). In tricaine-treated embryos, the number of serotonergic neurons appears normal. The shortened arms of tricaine-treated embryos result in crowding of the synB neurons; however, their number and spatial arrangement appears normal in that context (Fig. S6B). Thus, tricaine does not affect neural specification or patterning at 48 hpf. We also assessed the ciliary band (CB), which is a narrow ectodermal region at the boundary between the dorsal and ventral territories (Yaguchi et al., 2010). Since the CB fate is repressed by DV specifying TGFβ signals, it is a useful readout for DV perturbations (Bradham et al., 2009; Duboc et al., 2004; Yaguchi et al., 2010). We found that the CB is restricted to a normal, narrow band in both control and tricaine-treated embryos (Fig. S6C-D), indicating that tricaine treatment does not perturb DV specification. To corroborate that result, we performed single molecule (sm) FISH for LvChordin and LvIrxA, markers of ventral and dorsal, respectively. The spatial expression of each gene was not significantly affected by tricaine treatment relative to controls, nor was the inferred size of the CB between them (Fig. S6E-K), indicating that VGSC activity is not required for ectodermal DV specification.

### Skeletal patterning cues in the ventrolateral ectoderm are perturbed in tricaine-treated embryos

The posterior ventrolateral (VL) ectoderm corresponds with the expression domain for two ectodermal skeletal cues, LvWnt5 and LvVEGF, and is posterior to the expression of a third cue, Univin (Adomako-Ankomah and Ettensohn, 2013; Duloquin et al., 2007; McIntyre et al., 2013; Piacentino et al., 2015; Piacentino et al., 2016b). To test whether VGSC inhibition impacts the expression of these genes, we next performed quantitative smFISH (Choi et al., 2018; Choi et al., 2020) for each cue in control and tricaine-treated embryos (Fig. 4A-B). Focusing on the lateral expression territories where the expression of each signal is enriched, we found that both LvWnt5 and LvVEGF, but not LvUnivin, are significantly spatially expanded in tricaine-treated embryos relative to controls, with the most dramatic effect on Wnt5 (Fig. 4C1). We also quantified the expression level within the same regions, and similarly identified a significant increase in Wnt5 and VEGF, but not Univin expression (Fig. 4C2). To evaluate whether the increased levels reflect increased transcription or are instead mainly due to spatial expansion, we determined the average expression per unit area within each lateral territory and found that only Wnt5 expression exhibits a mild but significant increase; in contrast, the local VEGF and Univin expression levels were not changed by tricaine (Fig. 4C3). Thus, VGSC inhibition is sufficient to spatially expand and elevate the expression Wnt5 in the ventrolateral regions and to mildly spatially expand VEGF in the same spatial locations but is insufficient to affect the expression of the spatially adjacent anterolateral cue Univin.

**Figure 4.**
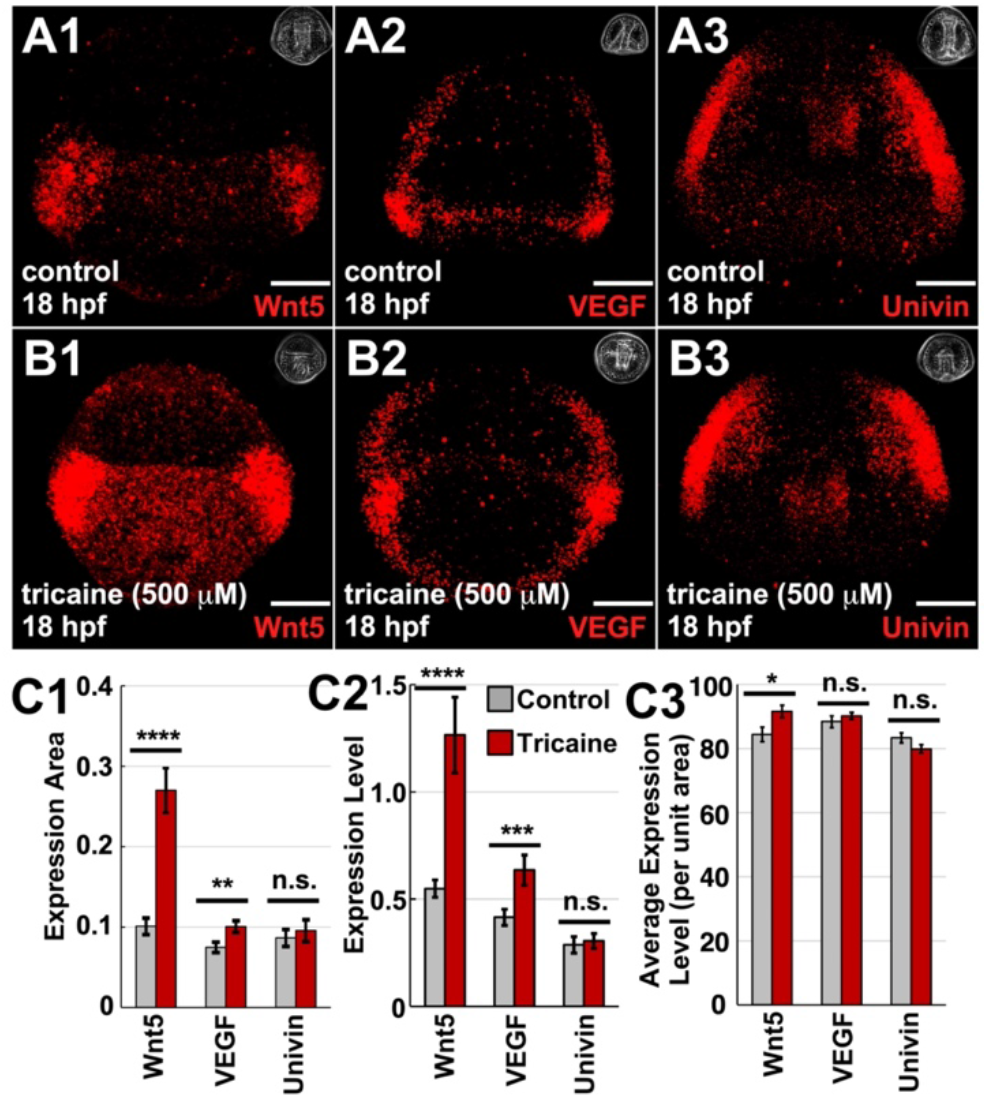
VGSC activity is required to spatially restrict the gene expression domains of Wnt5 and VEGF but not Univin. A-B. Control (A) and tricaine-treated (B1) late gastrula-stage embryos (18 hpf) were subjected to HCR FISH for the ectodermal patterning genes Wnt5 (1), VEGF (2), or Univin (3). C. The normalized area of the lateral expression domains (C1), the total normalized expression level within those domains (C2), and the normalized average expression per unit area (C3) for each gene is shown as the average ± s.e.m.; n ≥ 30 (Wnt5, VEGF) or n = 17 (Univin); * p < 0.05; ** p < 10^−2^; ***, p < 10^−3^; **** p < 10^−4^; n.s., not significant *t*-test). See Methods for details. Scale bars represent 20 *µ*m.

### VGSC activity is required for normal PMC migration and syncytial integrity

Since VGSC inhibition elicits skeletal patterning defects and perturbs the spatial expression of the ectodermal patterning genes VEGF and Wnt5, we next evaluated the effects of tricaine on the skeletogenic PMCs. In late gastrula-stage controls, the PMCs are arranged in the ring-and-cords pattern with ventrolateral clusters in which skeletogenesis initiates; the PMCs are connected by a syncytial cable in which the biomineral is deposited (Fig. 5A1). In tricaine-treated late gastrula-stage embryos, the PMCs approximate the ring-and-cords pattern; however, there are a number of anomalies (Fig. 5A4), including breaks in the syncytial cable, “rogue” PMCs that failed to incorporate into the syncytium, and ectopic PMC clusters. We quantified the frequency of breaks and ectopic clusters, and found that, at late gastrula stage, 100% of the tricaine-treated embryos exhibited a syncytial break and 75% exhibited at least one ectopic cluster (Fig. 5B). Syncytial breaks were most common in the ventral part of the ring and least common in the cords at late gastrula stage (18 hpf) and were fewer thereafter (Fig. S7A), implying that tricaine-treated embryos on average were able to resolve some of the PMC anomalies after late gastrula stage. Rogue PMCs were significantly increased among tricaine-treated embryos at each of the assessed time points (Fig. S7A).

**Figure 5.**
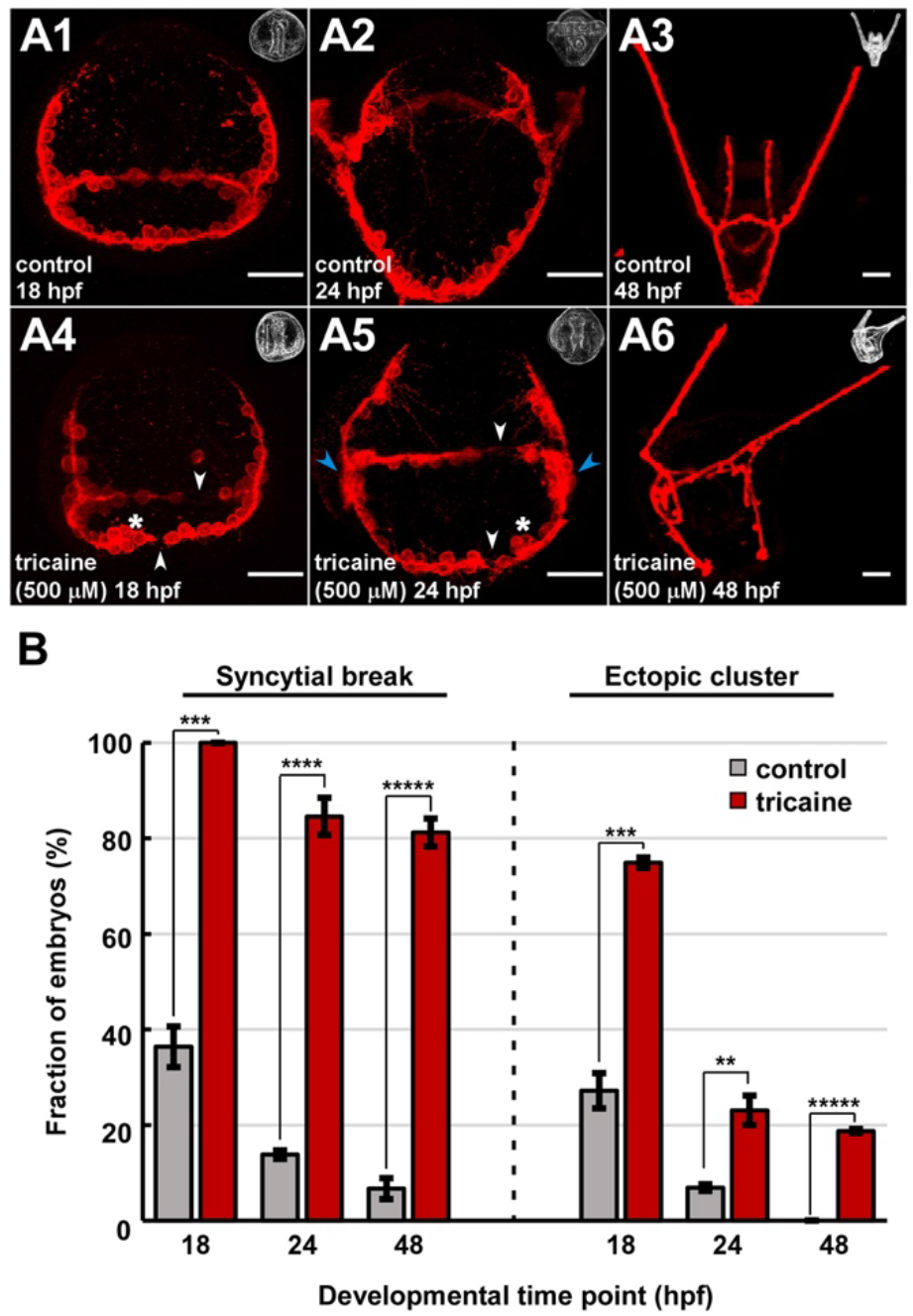
VGSC activity is required for normal PMC migration and syncytial integrity. A. Immunolabeled PMCs in control (1-3) and tricaine-treated (4-6) embryos are shown at the indicated timepoints; corresponding phase contrast images are inset. Defects include syncytial breaks (white arrow-heads), ectopic clusters (asterisks), and lack of normal migration into the secondary pattern (blue arrow-heads). B. The fraction of embryos exhibiting each defect is shown as the average percentage ± s.e.m.; n ≥19 for control and tricaine at 18 and 24 hpf; n ≥ 15 for control and tricaine at 48 hpf; * p < 0.05; ** p < 10^−4^; *** p < 10^−8^; **** p < 10^−12^; ***** p < 10^−16^; n.s. not significant (weighted *t*-test). See also Fig. S7. Scale bars in A1, 2, 4, and 5 are 20 *µ*m; the remainder are 50 *µ*m.

In 24 hpf control embryos (late prism stage, Fig. 5A2), the PMCs have begun migrating from the primary PMC pattern to form the secondary elements. However, in tricaine-treated 24 hpf embryos, the PMCs fail to begin secondary migration and appear stalled in the late gastrula-stage pattern (Fig. 5A5). In tricaine-treated late plutei, the PMCs have migrated to some but not all of the normal secondary positions; abnormal PMC positioning that corresponds with skeletal patterning defects is evident (Fig. 5A6). Additional PMC abnormalities persist but were diminished in tricaine-treated plutei, including syncytial breaks and ectopic PMC clusters (Fig. 5B, S7B). The large reduction in ectopic clusters after late gastrula stage might reflect the normal migration of PMCs from the clusters (including ectopic clusters), diminishing them, during secondary skeletal patterning. Consistent with that possibility, we observed extra triradiates in some tricaine-treated embryos that probably reflect these prior ectopic PMC clusters; the ectopic skeletal rods induced by tricaine might also reflect partial development of ectopic triradiates (Fig. 1B). Finally, while variable, the number of rogue PMCs is significantly increased in tricaine-treated embryos at each timepoint, and most dramatically at 48 hpf (Fig. S7A-B). We also note a surprising reduction in rogue PMCs in each condition at 24 hpf, suggesting that this population undergoes dynamic changes rather than remain-ing a static group that remains disconnected from the PMC syncytium. Taken together, these data show that VGSC activity is required for timely PMC migration, normal spatial positioning of PMCs, continuity of the PMC syncytium, prevention of rogue PMCs, and restriction to only two PMC clusters.

### VGSC activity is required to spatially restrict expression of c-Jun and Pks2

Over the course of skeletal patterning, the PMCs diversify, reflected by subpopulations of PMCs that exhibit differential gene expression; this diversification likely occurs in response to the receipt of patterning cues (Croce et al., 2003; Gross et al., 2003; Piacentino et al., 2016b; Sun and Ettensohn, 2014; Zuch and Bradham, 2019). We next investigated whether expression of three PMC subset genes, LvJun, LvPks2, and LvVEGFR, is perturbed by VGSC inhibition using smFISH and PMC immunostaining. Jun is a transcription factor that is expressed in the PMC clusters at late gastrula stage (Fig. 6A) (Sun and Ettensohn, 2014). Notably, the ectopic PMC clusters that arise in tricaine-treated late gastrula embryos strongly express Jun, while rogue PMCs typically do not (Fig. 6B, G). Pks2 is a PMC subset gene that encodes a protein which belongs to the group of polyketide synthases, and it is expressed in PMCs in the ventral and dorsal midline, clusters, and anterior tips of the cords at late gastrula stage (Fig. 6C) (Castoe et al., 2007; Sun and Ettensohn, 2014; Zuch and Bradham, 2019). In tricaine-treated embryos, Pks2 is expressed at highly elevated levels in the same PMC subsets as controls while also expanding into all of the PMCs in the ventral part of the PMC ring as well as the rogue PMCs (Fig. 6D, 6H). LvVEGFR expression is present throughout the PMC complement at late gastrula stage and is elevated in the cluster PMCs adjacent to the ectodermal sites of VEGF expression (Adomako-Ankomah and Ettensohn, 2013; Duloquin et al., 2007; Piacentino et al., 2016a; Piacentino et al., 2016b; Schatzberg et al., 2015; Sun and Ettensohn, 2014); this expression profile is unaffected by tricaine (Fig. 6E-F, 6I). Taken together, these data show that VGSC inhibition is sufficient to perturb PMC subpopulation identity as reflected by the abnormal spatial expression profiles for Jun and Pks2 in tricaine-treated embryos. Since Jun and Pks2 both exhibit spatially expanded expression, the results suggest that VGSC activity normally functions to restrict Jun and Pks2 expression spatially and to reduce Pks2 expression levels.

**Figure 6.**
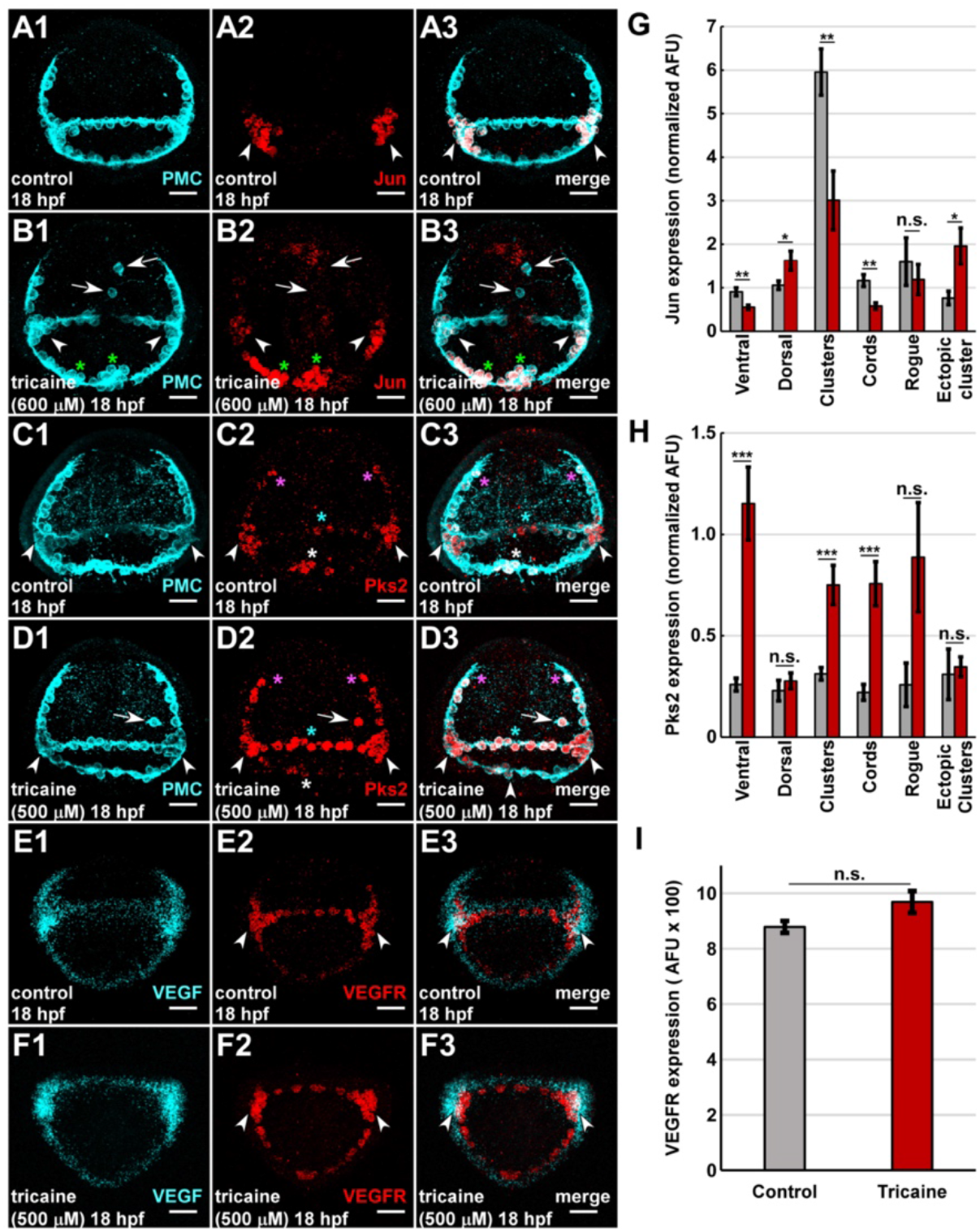
VGSC activity is required for normal spatial expression of PMC subset genes Jun and Pks2 at late gastrula stage. A-F. Control (A, C, E) and tricaine-treated (B, D, F) late gastrula-stage (18 hpf) embryos were subjected to HCR FISH for Jun, pks2, or VEGFR (red, 2) and PMC immunolabeling (cyan, A1, B1, C1, D1) or FISH for VEGF (cyan, E1, F1), and are shown as confocal z-projections. PMC clusters (arrowheads), rogue PMCs (arrows), the dorsal midline/sheitel (white asterisks), the ventral midline (cyan asterisks), the cord tips (pink asterisks), and ectopic clusters (green asterisks) are indicated. G-I. The expression levels for each gene are shown as the normalized average signal ± s.e.m. within the indicated PMC territories and within rogue PMCs; n ≥ 15; * p < 0.05; ** p < 10^−2^; *** p value < 10^−3^; n.s. not significant (*t*-test). Scale bars are 20 *µ*m.

### Wnt5 reduction is sufficient to rescue tricaine-mediated skeletal patterning defects

Since both Wnt5 and Scn5a are expressed by the posterior VL ectoderm, we next confirmed their spatial overlap using smFISH. smFISH for LvScn5a reveals a general background of expression in many or most cells, particularly in the anterior half of the embryo, along with higher levels of Scn5a expression in the ventrolateral and apical domains of the ectoderm at 18 hpf (Fig. 7A2), similar to but more sensitively than standard FISH (Fig. 3B, S4). These experiments confirm that tricaine treatment results in a dramatic expansion of the Wnt5 spatial expression domain at late gastrula stage, and that Wnt5 and Scn5a expression spatially overlap in the posterior ventrolateral ectoderm in both controls and in tricaine-treated embryos (Fig. 7A, arrow-heads). Tricaine treatment also resulted in generally elevated Scn5a gene expression (Fig. 7A5). These results indicate that LvScn5a is a house-keeping gene expressed by most or all cells, consistent with other models (Black and Waxman, 2013; de Lera Ruiz and Kraus, 2015; Kaestner et al., 2018; Roger et al., 2015). From these results along with similar data in Fig. 4, it seems likely that Scn5a normally exerts a globally repressive effect on Wnt5 expression, with relatively complete effects that prevent Wnt5 expression outside the VL ectoderm, along with incomplete effects that reduce the level and spatially restrict Wnt5 expression within the VL ectoderm; the difference within the VL ectoderm presumably results from a locally expressed Wnt5 transcriptional activator that counters the repressive effects that result from Scn5a activity.

**Figure 7.**
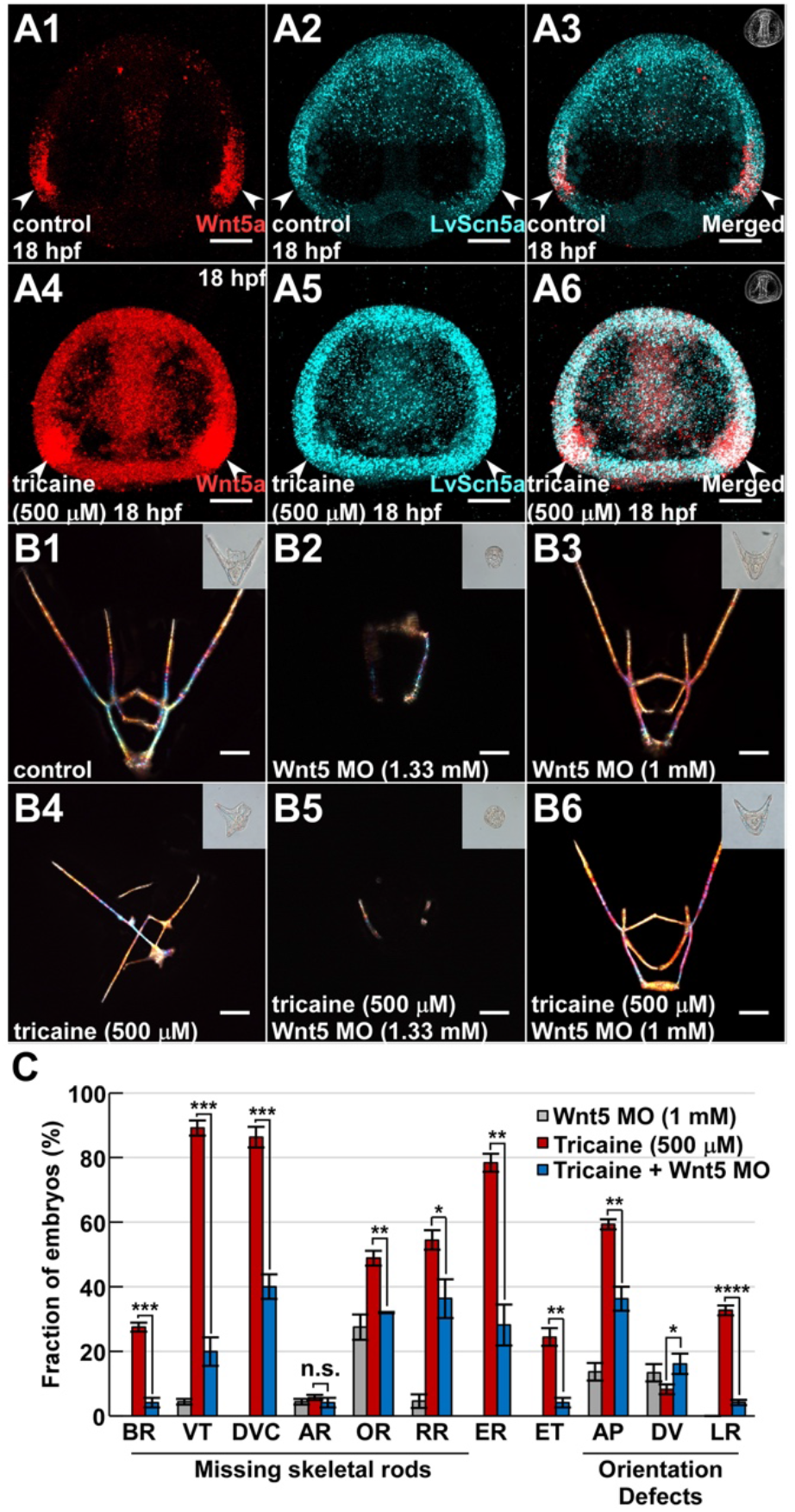
Suboptimal Wnt5 knockdown is sufficient to rescue the tricaine phenotype. A. Control (A1-3) and tricaine-treated (A4-6) embryos subjected to HCR FISH at late gastrula stage for Wnt5 (red) and LvScn5a (cyan) are shown as confocal z projections. The spatial overlap of expression of these genes in the posterior VL ectoderm is indicated (arrow-heads). B. Control embryos (B1, B4) and embryos microinjected with Wnt5 MO at a high dose (B2, 5) or low dose (B3, 6) are shown at the pluteus stage without (B1-3) or with tricaine treatment (B4-6) as skeletal birefringence images with corresponding DIC images inset. C. The frequency of the indicated skeletal patterning defects per embryo is shown as the average percentage ± s.e.m.; n ≥ 22 per condition; * p < 0.05; ** p < 10^−4^; *** p < 10^−8^; **** p < 10^−12^; ***** p < 10^−16^; n.s. not significant (weighted *t*-test). Scale bars in A represent 20 *µ*m; the remainder are 50 *µ*m.

We next asked whether the skeletal patterning defects associated with VGSC inhibition could be rescued by Wnt5 suppression. To test this, zygotes were microinjected with LvWnt5 MO and treated with tricaine, then assessed for skeletal patterning defects at the pluteus stage. Injection of Wnt5 MO at a high dose resulted in inhibition of skeletogenesis, with arrested development at late gastrula stage, consistent with previous findings (McIntyre et al., 2013) (Fig. 7B2, B5). Because our goal was to evaluate skeletal patterning, we performed a dose-response of Wnt5 MO and selected a suboptimal dose that permitted skeletogenesis and morphogenesis to occur. The majority of embryos injected with a suboptimal dose of Wnt5 MO developed control-like skeletons (Fig. 7B3, C). When Wnt5 MO injection was combined with tricaine treatment, skeletal patterning defects were significantly rescued (Fig. 7B6, C). These results indicate that Wnt5 suppression is sufficient to significantly rescue the skeletal patterning defects induced by VGSC inhibition. These findings suggest that the major skeletal patterning defects associated with tricaine treatment can be explained by the increase in Wnt5 expression level and spatial extent, including the AP orientation defects.

### VEGF reduction is not sufficient to rescue tricaine-mediated skeletal patterning defects

We did not observe a change in average VEGF expression level in the VL ectoderm in response to tricaine, although VEGF spatial expression was mildly expanded, and the total level of VEGF expression was correspondingly increased (Fig. 4). Previous studies indicated that Wnt5 activity negatively regulates VEGF expression (McIntyre et al., 2013), raising the question of whether VGSC inhibition ultimately impacts skeletal patterning through an indirect effect on VEGF via impacts on Wnt5. To test whether increased VEGF is functionally relevant for the tricaine phenotype, we treated embryos with the established VEGF inhibitor axitinib (Adomako-Ankomah and Ettensohn, 2013; Bhargava and Robinson, 2011; Piacentino et al., 2016b; Sun and Ettensohn, 2014). High doses of axitinib delivered prior to gastrulation block skeletal development, while treatment in Lv after 16 hpf is permissive to skeletogenesis but results in specific patterning defects (Adomako-Ankomah and Ettensohn, 2013; Piacentino et al., 2016b).

Since tricaine is effective prior to 16 hpf (Fig. S1), we treated embryos with both drugs from fertilization onward, using a range of suboptimal doses of axitinib that are permissive to skeletal development. Embryos exposed to tricaine along with axitinib did not exhibit a rescue of patterning defects at any dose (Fig. 8), indicating first that tricaine-mediated spatial expansion of VEGF does not account for the ensuing skeletal patterning defects, and second, that the phenotypic effects of Wnt5 are not mediated by VEGF.

**Figure 8.**
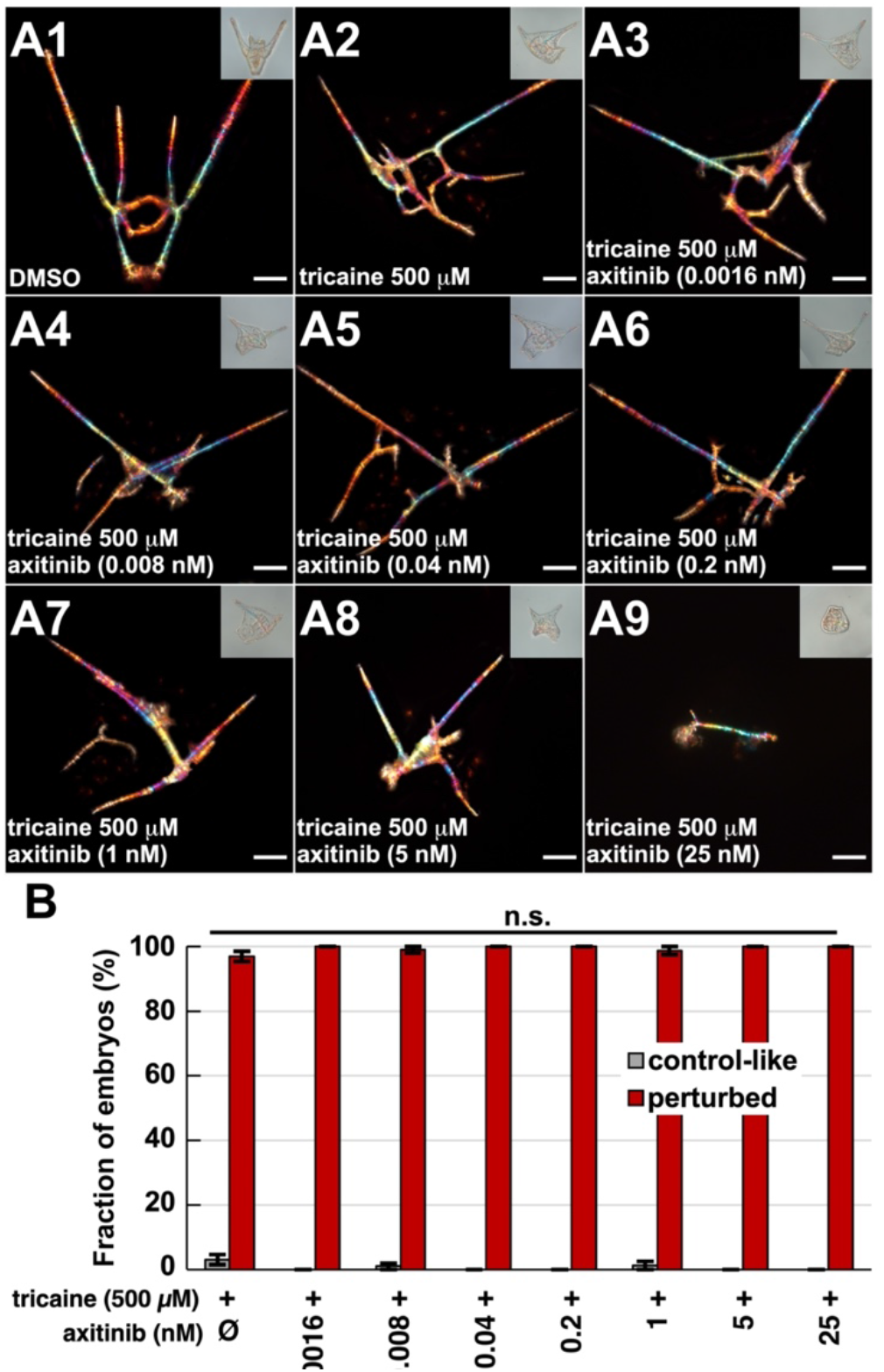
Inhibition of VEGF signaling is insufficient to rescue tricaine-induced skeletal patterning defects. A. Vehicle-treated (1), tricaine-treated (2), and axitinib and tricaine co-treated (3-9) embryos are shown at the pluteus stage as skeletal birefringence images with corresponding DIC images inset. Embryos were treated continuously with tricaine and/or axitinib from fertilization. B. The number of control-like and perturbed embryos per condition is shown as the average percentage across three biological replicates ± s.e.m.; n ≥ 112 per condition; n.s. not significant (t-test). Scale bars in A are 50 *µ*m.

### Model

These results are consistent with a model in which VGSC activity, both in the ventrolateral ectoderm and in general, functions to spatially restrict Wnt5 expression and signaling. In the chronic absence of VGSC activity, ventrolateral Na^+^ levels are restored to normal, presumably by compensatory effects, prior to the temporal period of sensitivity. During this sensitive window, Wnt5 expression becomes spatially expanded in the posterior ventrolateral ectoderm, ectopic Jun-expressing PMC clusters and triradiates arise, and abnormal skeletal patterning ensues (Fig. 9).

**Figure 9.**
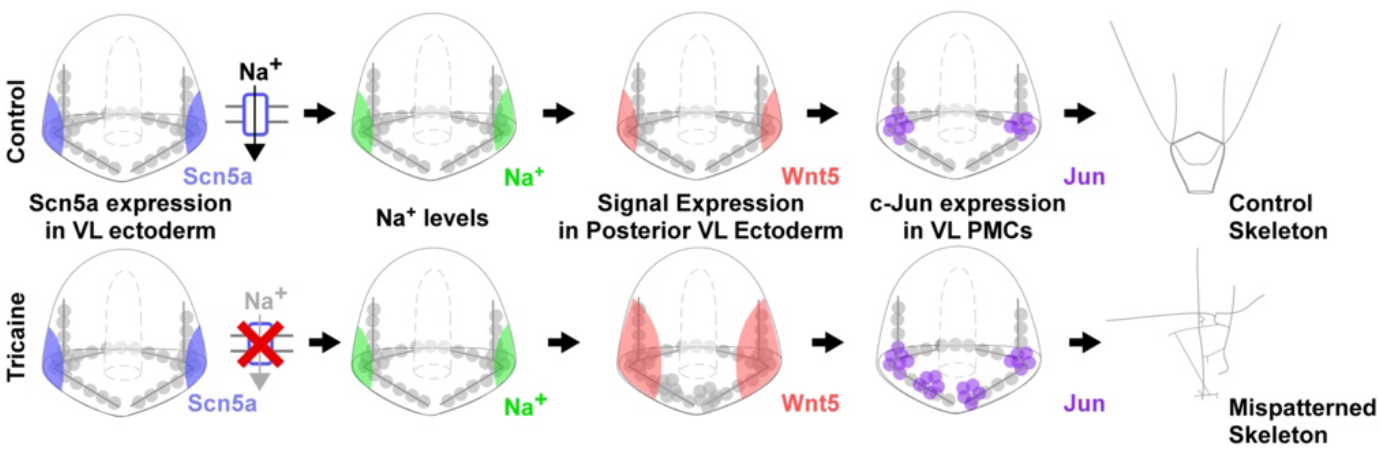
A model for tricaine-mediated skeletal patterning defects. In controls (upper panel), Scn5a expressed in the posterior VL ectoderm is required for spatially restricted Wnt5 expression in the same ectodermal regions. Adjacent to the VL ectoderm, Jun-expressing PMC clusters produce triradiates; normally patterned skeletons subsequently develop. Chronic VGSC inhibition (lower panel) results in Na^+^ levels restored to normal, spatially expanded Wnt5 expression, ectopic Jun-expressing PMC clusters, and abnormal skeletal patterns that include ectopic elements.

## Discussion

Overall, our study presents the novel finding that the activity of LvScn5a, a voltage-gated sodium channel (VGSC), is a previously unrecognized, required regulator of skeletal patterning that functions to restrict the spatial expression of patterning cues VEGF and Wnt5. In this study, we used the local anesthetic tricaine to show that VGSC activity is required for normal skeletal patterning in sea urchin embryos. Inhibition of VGSC activity is sufficient to induce ectopic PMC clusters and triradiates along with skeletal patterning defects. Lv embryos express an endogenous VGSC gene, LvScn5a, in posterior ventrolateral (VL) ectodermal regions that are adjacent to the PMC clusters where skeletogenesis initiates. We show that tricaine-mediated skeletal patterning defects can be rescued by overexpression of an anesthetic insensitive version of LvScn5a, indicating the specificity of tricaine. Finally, we show that the tricaine-mediated skeletal patterning defects are rescued by reduction of Wnt5 expression, demonstrating that the expanded expression of Wnt5 is the functionally relevant effect of VGSC inhibition.

We show that acute tricaine treatment results in depressed Na^+^ levels and hyperpolarization, as expected. However, longer-term tricaine treatment results in both depolarization and a return of Na^+^ to levels that are comparable to controls. We therefore interpret the depolarization of long-term tricaine-treated embryos to be a result of compensatory mechanisms, likely via other ion channels that restore normal Na^+^ levels at the expense of V_mem_. Similar compensatory changes have been observed after prolonged inhibition of other ion channels (Schatzberg et al., 2015). Interestingly, tricaine-treated embryos appear to upregulate LvScn5a expression in response to tricaine treatment, which may also indicate an additional compensatory response.

VGSC inhibition results in the formation of ectopic PMC clusters and supernumerary skeletal triradiates. To date, VEGF is the only patterning cue known to be required for PMC cluster formation (Adomako-Ankomah and Ettensohn, 2013; Duloquin et al., 2007). Our results indicate that Wnt5 signaling is also important for PMC cluster formation. We find that Wnt5 expression in the posterior VL ectoderm is dramatically spatially expanded in response to VGSC inhibition. That expansion correlates with the formation of ectopic PMC clusters and triradiates as well as the production of ectopic skeletal elements, and importantly, is rescued by partial Wnt5 knockdown. That rescue suggests that Wnt5 expression in the VL ectoderm regulates PMC cluster formation and triradiate secretion, and thus identifies Wnt5 as a second cue, along with VEGF, that organizes PMC clusters.

Previous work showed that global overexpression of Wnt5 resulted in abnormal PMC organization (McIntyre et al., 2013). This is similar to our findings that local expansion of Wnt5 is sufficient to pattern ectopic PMC clusters. The prior study further suggested that Wnt5 is a repressor of VEGF expression in the posterior VL ectoderm (McIntyre et al., 2013). This finding might be interpreted as indicating that Wnt5’s effects on skeletal patterning are mediated entirely by VEGF. However, it is difficult to explain the spatially inappropriate positioning of PMCs in anterior regions in Wnt5-over-expressing embryos as entirely reflective of VEGF repression since the PMCs occupy anterior regions in those embryos where VEGF is already normally absent (McIntyre et al., 2013). Instead, it is more reasonable to posit that Wnt5 mediates PMC-organizing effects independently of VEGF, both in the previous work (McIntyre et al., 2013) and herein; thus, both results implicate Wnt5 as a direct signal that spatially organizes the PMCs.

We observed a mild spatial expansion of VEGF expression in tricaine-treated embryos that is not reciprocal to the dramatic expansion of Wnt5 expression and is therefore not consistent with Wnt5 acting as a VEGF repressor. Quantification of our results indicates that increased Wnt5 signal levels are not sufficient to repress VEGF expression levels as previously suggested (McIntyre et al., 2013), and indicate instead that the expression of VEGF and Wnt5 are at least partially independently regulated. Moreover, we show that inhibition of Wnt5, but not VEGF, is sufficient to rescue skeletal patterning defects, importantly indicating that Wnt5 signals independently of VEGF to mediate skeletal patterning.

It could be argued that the ectopic PMC aggregates observed in tricaine-treated embryos do not necessarily have PMC cluster identity and that instead, VGSC inhibition modulates cell adhesion, thereby producing these extra PMC aggregates. However, previous work showed that c-Jun expression marks the PMC clusters at late gastrula stage (Sun and Ettensohn, 2014), suggesting that tricaine-induced PMC aggregates are, indeed, ectopic PMC clusters. Moreover, the occurrence of ectopic, supernumerary triradiates in VGSC-inhibited embryos reinforces that conclusion. Our data are thus consistent with a model in which Wnt5 signaling from the posterior VL ectoderm is sufficient to organize ectopic clusters of PMCs that express c-Jun and produce triradiates or other skeletal rods. It remains unclear whether Wnt5 signals via canonical or non-canonical pathways in this context; however, each of the known Wnt5 pathways impacts gene expression, and could account for Wnt5-mediated activation of *jun* expression. This model suggests a novel and as of yet unreported role for Wnt5 in sea urchin skeletal patterning and prompts further inquiry into the relationship between Wnt5 signaling and PMC subset gene expression.

Our results suggest that VGSC activity exerts repressive control over Wnt5 expression, with the strongest effect on spatial regulation. We show that temporally, VGSC activity is required for normal skeletal patterning between 14-17 hpf, corresponding to early and mid-gastrula stages. We find that Wnt5 expression is spatially expanded at 18 hpf, immediately following the sensitive window. This is consistent with VGSC-dependent Wnt5 expression control occurring during the window of sensitivity to tricaine. Scn5a expression persists, particularly in neurogenic ectodermal regions, after 18 hpf, but no longer impacts skeletal patterning. It seems likely that the subsequent role of Scn5a is restricted to neural functions.

It remains mechanistically unclear how VGSC inhibition modulates Wnt5 expression; however, an impact of this channel on gene expression has also been observed in the context of metastatic cells (House et al., 2010), indicating that this is not a sea urchin-specific phenomenon. Tricaine treatment abolishes the regenerative response of the tail in *Xenopus*. This effect is due to changes in intracellular Na^+^ levels and is mediated by the salt inducible kinase (SIK) (Sanz, 2003; Tseng et al., 2010). Thus, SIK seems to be an intriguing prospect as a mediator of the Scn5a activity in sea urchins. However, LvSIK is expressed at low levels during skeletal patterning (Hogan et al., 2020). Further, in tricaine-treated embryos, we observe normal Na^+^ levels during the temporal window of VGSC functional relevance along with abnormal V_mem_, which is not consistent with effects mediated by perturbation to Na^+^ levels or to SIK.

An alternative possibility is that Scn5a activity negatively regulates Wnt5 spatial expression by interrupting a “community effect” by which Wnt5 expression spreads among cells via Wnt5 signaling that positively feeds back on Wnt5 expression. Such community effects regulate the expression of Wnt8 via β-catenin/TCF (Minokawa et al., 2005). While it is currently unknown whether Wnt5 similarly signals via β-catenin, Wnt5 signaling is sufficient to mediate gene expression changes (McIntyre et al., 2013), implying that a community effect could occur whether it relies on β-catenin/TCF or on a different transcriptional regulator. It is unknown how boundaries are established that halt community effects; our results are consistent with a model in which V_mem_ status negatively regulates this community effect. Specifically, it suggests that a mildly depolarized state is inhibitory, whereas the strongly depolarized state that results from VGSC inhibition is not, allowing the Wnt5 to be generally expressed along with allowing the VL expression domain to be abnormally large. If correct, this model would intriguingly tie biophysical states to gene expression control. In a correlate model, the extent of Wnt5 extracellular diffusion, transport, or receptor binding could be modulated by V_mem_ status to achieve a similar result by impinging on the community effect at the level of the signal.

A third possibility is that VGSC activity influences the expression of a microRNA (miR), in keeping with the role that miR-31 exerts to spatially modulate the expression of Wnt1 and Eve expression within the posterior ectoderm (Sampilo et al., 2021). While the action of a miR is unlikely to be solely responsible for the dramatic differences in Wnt5 expression mediated by tricaine, a globally expressed miR might contribute to normal general Wnt5 repression and could in turn be regulated by VGSC activity. However, while miRNAs are known to regulate ion channel expression (Gross and Tiwari, 2018; Gross et al., 2016; Liu et al., 2016; Sakai et al., 2017; Shao et al., 2016), evidence for the converse regulation is currently more sparse. One study demonstrates that the anion channel CFTR regulates miRNA expression in mammalian embryos via bicarbonate influx and subsequent downstream signaling events (Lu et al., 2012), while another study shows that epithelial sodium channels negatively regulate the expression of two miRNAs in mammalian embryos (Sun et al., 2014). Potential VGSC-mediated impacts on miRNA expression or on a Wnt5 community effect are not mutually exclusive: both might contribute to negatively regulating Wnt5 expression. A molecular understanding of how VGSC activity controls the spatial expression of Wnt5 will be an important next step in unraveling the mechanisms that regulate sea urchin skeletal patterning.

## Methods

### Reagents

Chemicals were obtained from Sigma Aldrich or Fisher Scientific. Ethyl 3-aminobenzoate methanesulfonate (tricaine) was obtained from Sigma Aldrich (St. Louis, MO), Bis-(1,3-diethylthiobarbituric acid) trimethine oxonol (DiSBAC, relative polarization) and CoroNa green, acetoxymethyl ester (CoroNa, sodium ions) were purchased from Invitrogen (Waltham, MA).

### Embryo culture, microinjection, drug treatments

Lytechinus variegatus adults were obtained from either Reeftopia (Miami, FL) or the Duke University Marine Labs (Beaufort, NC). Gamete harvesting, embryo culturing, in vitro transcription, and microinjections were performed as previously described (Bradham and McClay, 2006; Piacentino et al., 2015). The Wnt5 MO was obtained from GeneTools. The Wnt-5 MO sequence is: (5′-CGCTGGCAGACAAAGGGCGACTCGA-3′) (McIntyre et al., 2013). Dose-response experiments were performed with all reagents to determine their optimal working concentrations. Tricaine methanesulfonate (Sigma) was resuspended in artificial sea water (ASW) and was stored at −20°C in single use aliquots. Tricaine treatments were performed using artificial sea water (ASW) that was buffered to pH 8.1 using NaHCO_3_; corresponding controls were treated with the same buffered ASW vehicle. For acute tricaine treatments, embryos were cultured in ASW vehicle without inhibitor until time of imaging; they were then placed in a tricaine bath and imaged within 10 minutes.

### Morphological imaging and skeletal scoring

Embryos were imaged on a Zeiss Axioplan microscope at 200X with differential interference contrast (DIC) for morphology or with plane-polarized light to capture skeletal birefringence as a series of images in multiple focal planes. Montages of the focal planes were manually assembled using CanvasX (Canvas GFX, inc, Boston, MA) to present the entire skeleton in focus. All focal planes were used for scoring with our in-house scoring rubric (Piacentino et al., 2016), which encompasses element shortening, lengthening, loss, duplication, or transformation, ectopic element production, abnormal element orientation, as well as whole embryo-level defects such as midline defects. Left and right elements are separately scored. Elements are considered to be a loss when completely absent versus shortened when present but abnormally short (e.g., half their typical length or less). Ectopic elements occupy abnormal positions and do not correspond to a normal element (e.g., via their branching pattern). Orientation defects about each axis score the large-scale relationships between the skeletal components and are illustrated in Fig. 1 and in our prior work (Piacentino et al., 2016b; Rodriguez-Sastre et al., 2023). Briefly, AP defects reflect abnormally wide or narrow angles between left and right ARs (and BRs), DV defects are reflected by DVCs that are not parallel to each other or perpendicular to the AP axis, and LR defects are indicated by ARs that are oppositely directed. Some embryos exhibit multiple orientation defects. The landmarks for orientation analysis are the gut termini and the apical region of each larva. Midline defects lack any midline skeletal structures. Each of these defects is separately scored from individual embryos via manual inspection of montaged skeletal images along with individual focal planes. In some cases, skeletons were independently scored by different individuals to ensure objectivity. Control embryos exhibit patterning defect frequencies ranging from 0.5-1% for primary defects, and 3-10% for secondary and orientation defects. Orientation, midline, ectopic rods and ectopic triradiates are extremely rare among controls. Combined skeletal scores from seven experiments were subsampled into three equivalently sized bins for statistical analyses.

In some experiments in which large numbers of larvae were captured together in one focal plane at 50X, skeletons were simply scored as control-like or perturbed. In other similar experiments, larvae were scored in broad bins as exhibiting primary patterning defects (i.e., in the VTs, BRs, or DVCs), secondary patterning defects (i.e., in the ARs, ORs, or RRs), orientation defects (AP, DV, LR defects or any combination thereof), DV defects (radialization or notable ventral expansion), midline defects (completely separated left and right halves), uniform stunting (all elements short but otherwise normal), or toxicity (absence of gastrulation or morphogenesis). In these cases, ambiguous larvae (i.e., with overlapping or obscured elements) were scored conservatively as control-like.

### LvScn5a DR construct

An LvSnc5a sequence with the drug-resistant (DR) F1715A mutation was cloned into the pCS2 vector for *in vitro* transcription (Genscript, Inc. Piscataway, NJ).

### Fluorescent in situ hybridization (FISH)

Antisense riboprobes for LvVEGF, LvWnt5, and LvUnivin were previously described (Bradham et al., 2009; McIntyre et al., 2013; Piacentino et al., 2015; Schatzberg et al., 2021). A riboprobe corresponding to the open reading frame for LvScn5a was similarly produced. In situ hybridization was performed as previously described using digoxigenin-labeled probes (Piacentino et al., 2015).

### HCR FISH, Immunolabeling, and confocal microscopy

Hybridization chain-reaction (HCR) single molecule (sm) FISH probe sets were designed from the open reading frames for LvVEGF, LvWnt5, LvUnivin, LvScn5a, LvVEGFR, LvJun, LvPks2, LvChd, and LvIrxA by Molecular Instruments, Inc. (Los Angeles, CA, USA). Embryos were fixed in 4% paraformaldehyde then subjected to HCR smFISH (Choi et al., 2016; Choi et al., 2018) using fluorescently labeled amplifiers, buffers, and probe sets from Molecular Instruments, Inc. per the manufacturer’s instructions. Embryos were incubated in hairpin solution overnight in the dark at room temperature, washed with 5X SSCT, then mounted in PBS with 50% glycerol for imaging.

Immunofluorescent labeling was performed as previously described (Bradham et al., 2009). Primary antibodies were neural-specific monoclonal 1e11 (1:10, a gift from Robert Burke, University of Victoria, BC, Canada), anti-serotonin polyclonal (1:1000, Sigma-Aldrich), ciliary band-specific monoclonal 295 (undiluted; a gift from David McClay, Duke University, Durham, NC), and PMC-specific monoclonal 6a9 (1:5; a gift from Charles Ettensohn, Carnegie Mellon University, Pittsburgh, PA). Fluorescent secondary antibodies Cy2-conjugated goat anti-rabbit (1:900) and Cy3-conjugated goat anti-mouse (1:300) were obtained from Jackson Labs (West Grove, PA).

Confocal imaging was performed using an Olympus FV10i laser-scanning confocal microscope. For each experiment, all microscope settings were optimized on controls, then maintained across the experiment to ensure comparability. Confocal z-stacks were maximally projected using Fiji, and maximum intensity z-projections are presented.

### Fluorescent ion reporters

CoroNa and DiSBAC were used as previously described (Rodriguez-Sastre et al., 2019). Embryos were imaged live on an Olympus FV10i laser scanning confocal microscope, and, since these dyes rapidly photobleach, only single z-slices from the approximate center of each embryo were collected.

### Quantification

In all cases, quantifications are averages from ≥ three biological replicates unless otherwise noted. Statistical significance was determined with paired, two-tailed student *t-*tests for pair-wise comparisons; for comparisons among larger groups, the Tukey HSD test (Tukey, 1949) was employed. Identical acquisition settings were used across all embryos within each experiment to ensure their comparability.

#### PMC spatial scoring

Using confocal image stacks from PMC labeling experiments, PMCs from different regions of the overall pattern were manually subgrouped and counted. PMCs within the ventrolateral clusters were defined as contacting more than two other PMCs; the clusters delimit and segregate the ventral and dorsal parts of the PMC ring and the PMC cords.

#### PMC subset gene expression levels

For LvJun and LvPks2 smFISH results, ROIs were manually drawn around each PMC using Fiji, and the mean fluorescence intensity of the Jun or Pks2 signal was measured and normalized to z-depth by dividing by either the Hoechst (nuclei) or PMC stain fluorescence intensity for the respective PMC. Each image was background corrected by sampling empty space at the corners of the images then subtracting the average background per area from other signal intensities.

#### Ectodermal genes and fluorescent reporters

To quantify CoroNa, DiSBAC, and HCR FISH images, raw 16-bit images of individual z-slices were analyzed using Fiji by placing ROIs in distinct locations throughout the embryo in a stereotypic manner across all embryos. Within each ROI, the summed integrated density was used to measure the expression level, and the mean fluorescence value per unit area was also measured from each ROI to assess changes in gene expression level. Each image was background corrected as described above. The results were normalized to the maximum measured control value per condition as unitless percentages, then averaged.

#### Spatial gene expression analysis

For LvWnt5, LvVEGF, and LvUnivin smFISH results, to objectively determine the size of the enriched lateral expression domains, ROIs were assigned to them in an automated manner as follows. First, to focus specifically on the enriched region, a threshold was applied to z-projected embryos in Fiji to produce binary images. ROIs outlining the binarized expression domains were then defined in an automated manner using the Analyze Particles function in Fiji. Thresholding parameters were manually adjusted for each experiment to most accurately capture the enriched expression by optimizing the settings on representative control embryos, then applying those settings uniformly across the group. The resulting area values were then normalized to the total area of each z-projected embryo, rendering each measure unit-less.

For LvChd and LvIrxA FISH, embryos were imaged in vegetal views using an FV10i laser-scanning confocal microscope to allow for clear discrimination between ventral and dorsal territories. Confocal z-stack projections were analyzed in CanvasX (Canvas GFX, Inc., Boston, MA). Using the gut as the origin, the angles of the LvIrxA and LvChd expression domains were measured. The size of the inferred ciliary band was determined per embryo by subtracting the sum of the Chd and IrxA angles from 360°.

## Supporting information

Thomas et al Supplemental Material

## Author Contributions

This study was designed by CAB and CFT. The experiments were executed by CFT, VS, SM, ZY, and JG; data analyses were performed by CFT, DYH, VS, SM, ZY, JG, and CAB. The manuscript was written by CFT and CAB and edited by all co-authors.

## Acknowledgments

We thank Professors Charles Ettensohn, Robert Burke, and David McClay for antibodies and Dr. Todd Blute for microscopy advice. This work was funded by NSF IOS 1656752 (CAB). VS was partially supported by the BU Undergraduate Research Opportunities Program (UROP) and SM and JG by the BU Biology Summer Undergraduate Research Fellowship (SURF) program.

