## Supplementary material for "Voltage-gated sodium channel activity mediates sea urchin larval skeletal patterning through spatial regulation of Wnt5 expression": Thomas et al Supplemental Material

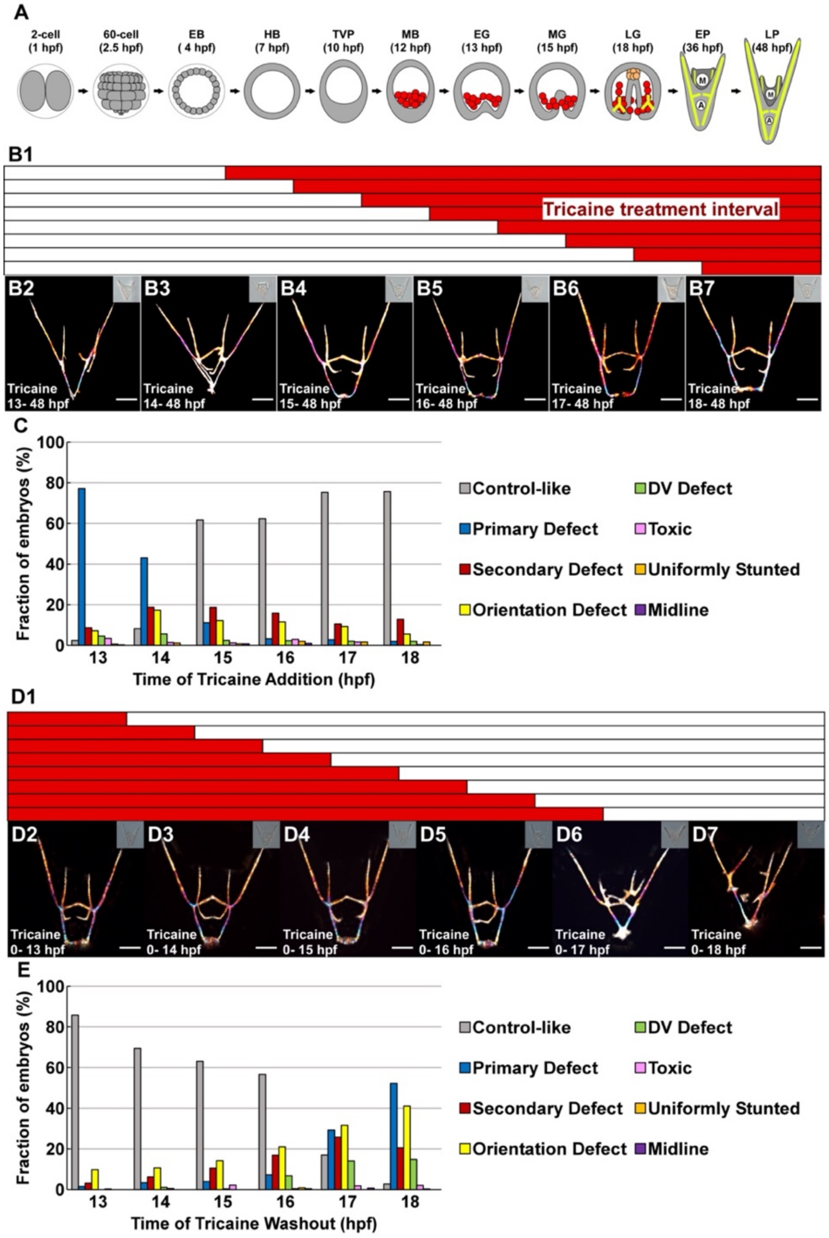


**Supplemental Figure 1. The frequency of skeletal patterning defects depends on the temporal interval of tricaine exposure.**

A. Sea urchin developmental stages are shown schematically. B. The schematic illustrates the experimental design for timed tricaine addition (red bars, B1). Corresponding exemplar embryos treated with tricaine during the indicated intervals are shown as skeletal birefringence images with corresponding DIC images inset (B2-7). C. Patterning defects during each interval of tricaine exposure are shown as the fraction of embryos that exhibited the indicated defects; n ≥ 247 per condition. D. The schematic illustrates the experimental design for timed tricaine removals (red bars, D1). Corresponding exemplar embryos are shown as in B (D2-7) and scored as in C (E); n ≥ 177 per condition. A. was adapted from Hogan, et al., 2020. See also Fig. 1.

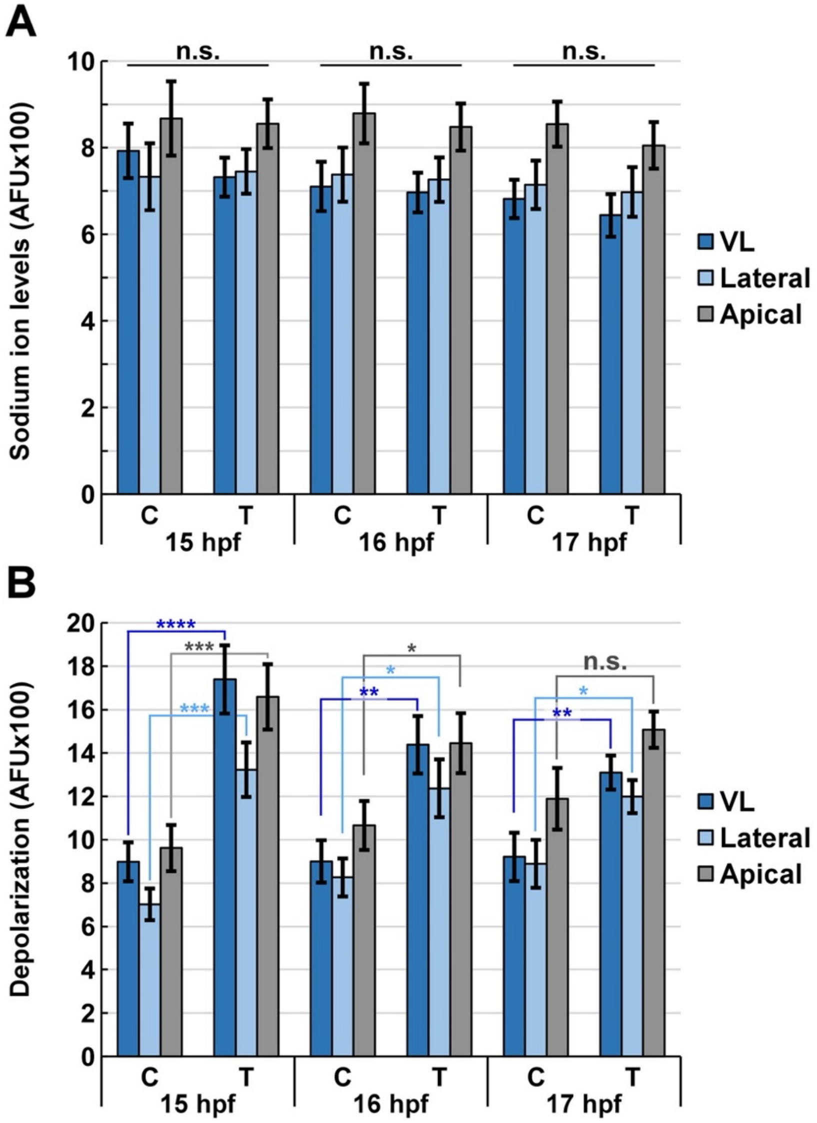


**Supplemental Figure 2. Long-term VGSC inhibition is sufficient to perturb voltage ($V_{mem}$) but not intracellular sodium ion levels.**

Sodium ions were visualized with the fluorescent reporter CoroNa (A), while $V_{mem}$ was visualized with DiSBAC in the same live embryos at the indicated time points (B). The results are shown as the mean AFU ± s.e.m.; n = 24 per condition; * $p < 0.05$; ** $p < 10^{-2}$; *** $p < 10^{-3}$; **** $p < 10^{-4}$; n.s. (student *t*-test). See also Fig. 2.

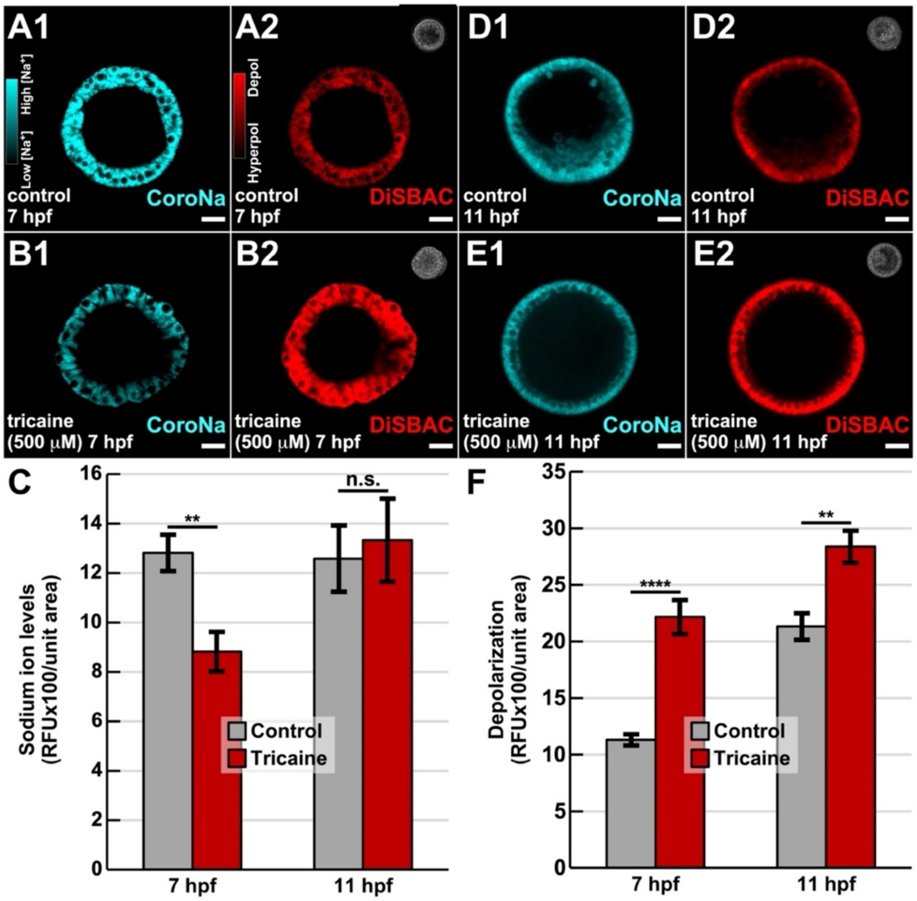


**Supplemental Figure 3. Prolonged VGSC inhibition results in compensatory changes to sodium ion levels and $V_{mem}$.**

A-B. Sodium ions were visualized with the fluorescent reporter CoroNa (cyan), and $V_{mem}$ was visualized with DiSBAC (red) together in live control embryos (A, D) and embryos treated with tricaine (B, E) from fertilization until either 7 (A-C) or 11 (D-F) hpf; the signal ranges are pseudocolored using custom monochrome LUTs (inset in A1, A2). The quantified results are shown as the average RFU/area ± s.e.m. (see also Fig. 2); n ≥ 20 embryos per condition; ** p < 10-3; **** p < 10-5; n.s. not significant (student t-test). Scale bars represent 20 μm.

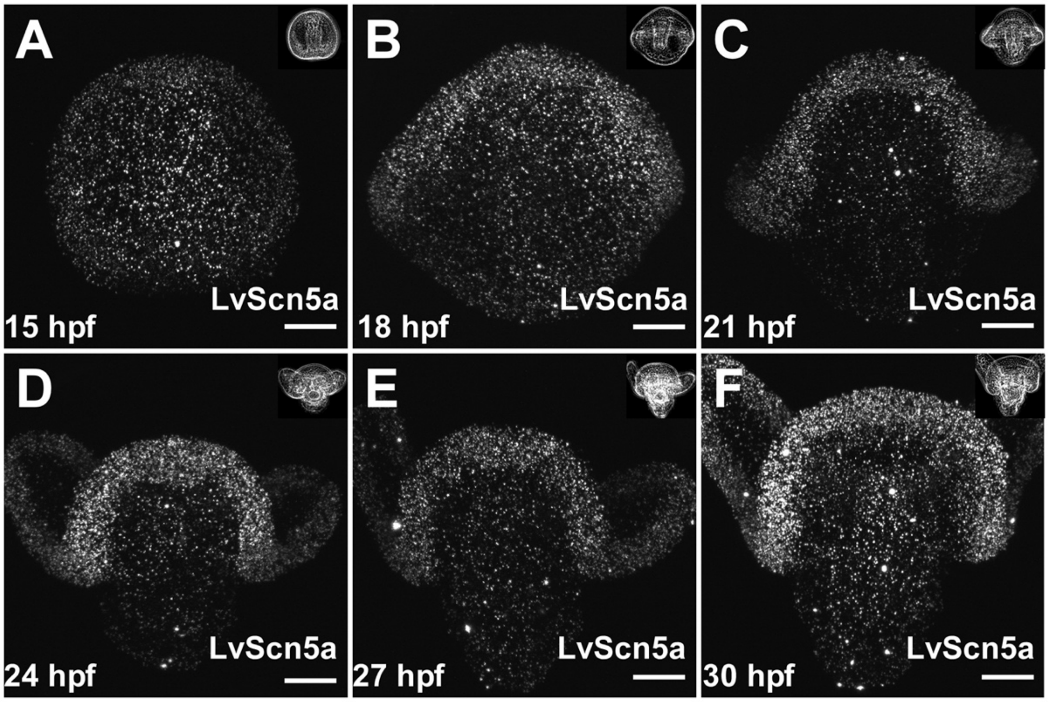


**Supplemental Figure 4. LvScn5a is broadly expressed at low levels and becomes enriched in the ciliary band as development proceeds.**
A-F. Control embryos were fixed at the indicated developmental timepoints, then subjected to FISH for LvScn5a. Raw LvScn5a expression is shown in z-projected embryos. See also Fig. 3B for the comparable background corrected, pseudocolored images.

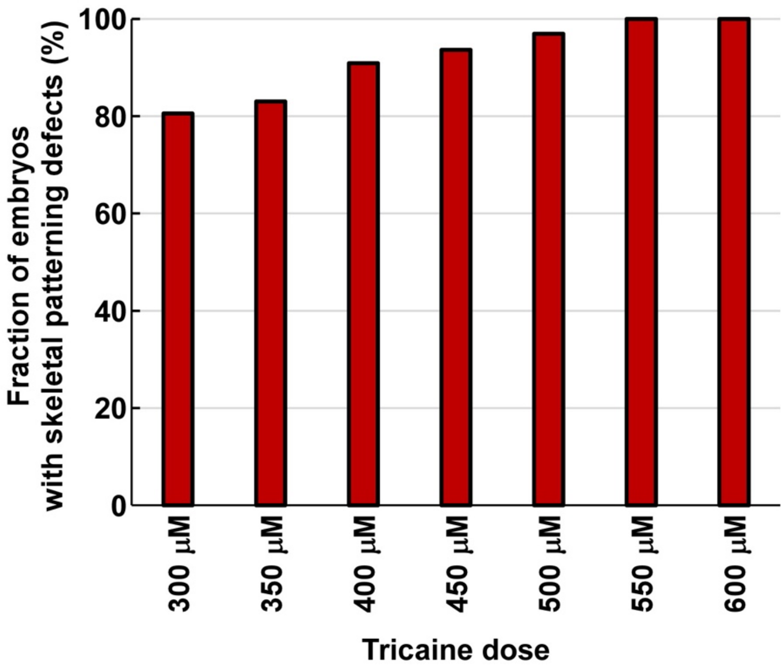


**Supplemental Figure 5. The fraction of embryos exhibiting skeletal patterning defects increases with dose.**
Embryos were exposed to the indicated doses of tricaine from fertilization to the pluteus stage, then examined to determine the percentage displaying patterning defects. n ≥ 37 for all conditions.

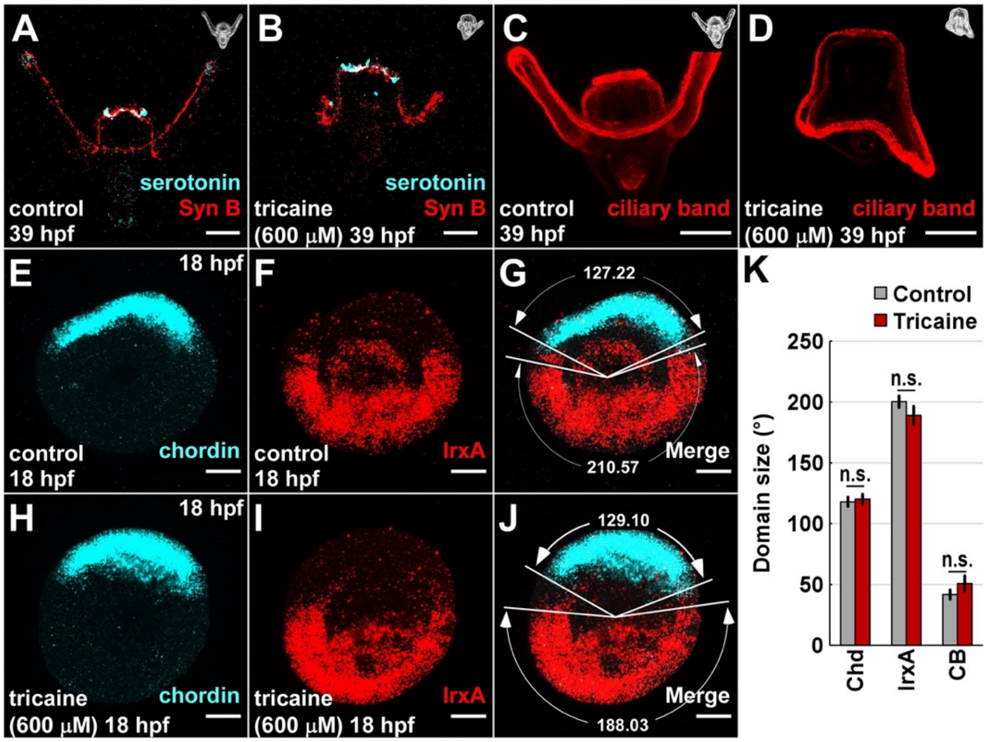


**Supplemental Figure 6. VGSC activity is not required for neural differentiation or DV specification.**

A-D. Control (A, C) and tricaine-treated (B, D) plutei (48 hpf) were immunolabeled for neurons as indicated (A, B) or for the ciliary band (C, D) and are shown as confocal z-projections with corresponding brightfield images inset. E-K. Control (E-G) and tricaine-treated (H-J) late gastrula-stage embryos subjected to HCR FISH for ventral chordin (cyan) and dorsal IrxA (red) are shown as confocal z-projections. The radial size of each of each expression domain was measured as shown (G, J); the size of the ciliary band (CB) was then inferred. The ventral (Chd), dorsal (IrxA) and ciliary band sizes are shown as the average angular degree ± s.e.m. per embryos; n ≥ 22; n.s., not significant (student t-test). (K). Scale bars are 20 $\mu$m.

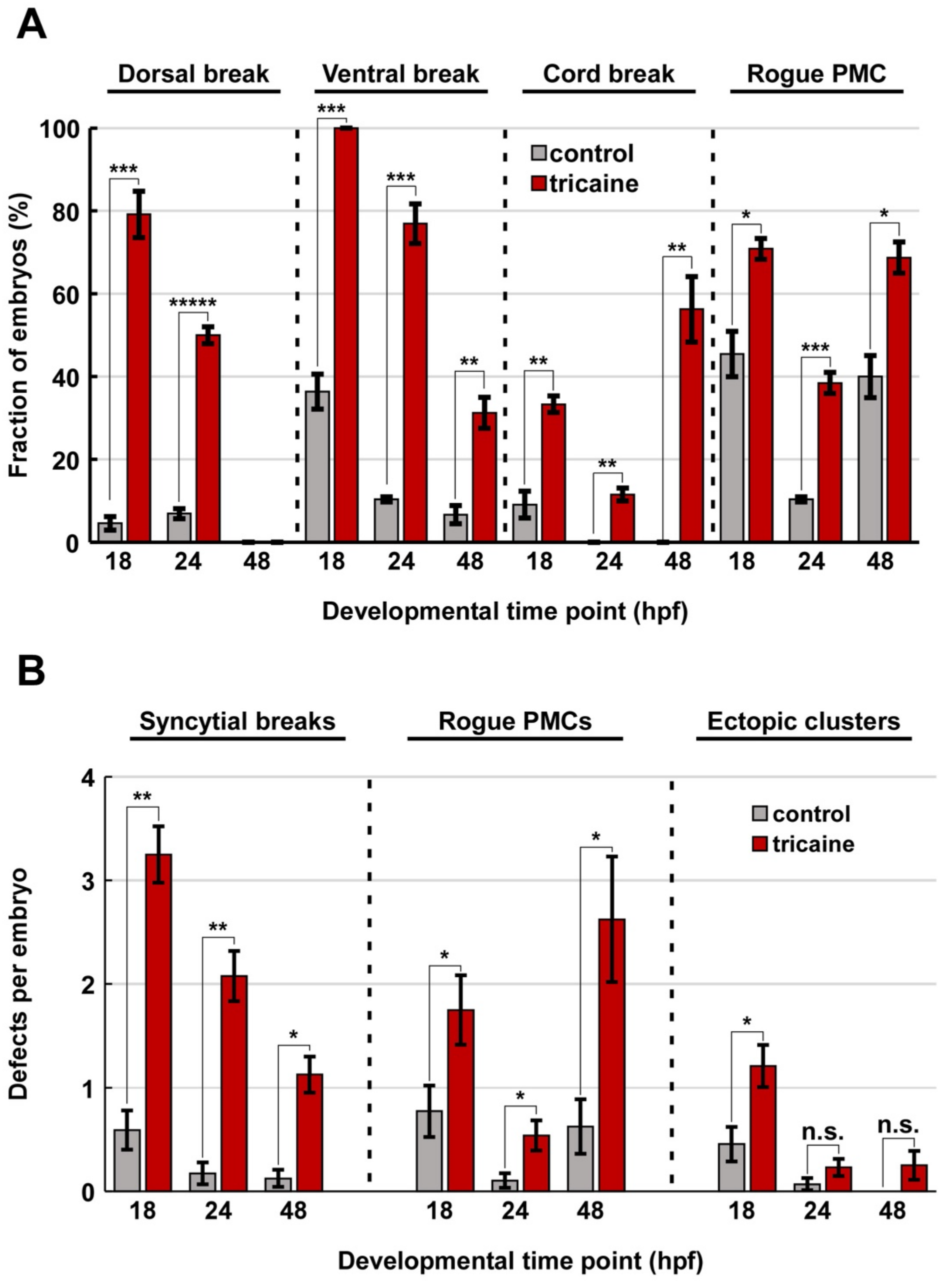


**Supplemental Figure 7. VGSC inhibition is sufficient to perturb PMC migration and syncytial integrity.**

PMC defects are shown as the percentage of embryos with each defect (A) or the number of defects per embryo per condition (B); data are shown as the average ± s.e.m.; n ≥ 15 per condition; * $p < 0.05$; ** p < 10-5; *** p < 10-10; **** p < 10-15; ***** p < 10-20; n.s., not significant (weighted t-test). See also Fig. 5.
